# A Tissue Virus Microenvironment with Activated Stress Responses Underlies Durable SIV Persistence

**DOI:** 10.64898/2026.04.02.716177

**Authors:** Thomas J. Hope, Eliana U. Crentsil, Muhammad S. Arif, Yanique Thomas, Ellen Zhang, Christopher T. Thuruthiyil, Ryan V. Moriarty, Flora Engelmann, Sean C. Pascoe, Jenna M. Hasson, Seth H. Borrowman, Maryam A. Shaaban, Edward J. Allen, Anne Monette, Ann M. Carias, Douglas Ferrell, Nawal Ouguirti, Isabelle Clerc, Judd F. Hultquist, Richard T. D’Aquila, Michael D. McRaven, Mariluz Araínga, Francois Villinger, Ramon Lorenzo-Redondo

## Abstract

HIV persistence during suppressive antiretroviral therapy (ART) remains a central barrier to cure, with the majority of reservoirs residing in gut-associated lymphoid tissues (GALT). Here, we define a spatially organized viral microenvironment (VME) that sustains reservoir durability and governs early viral rebound by comparing animals initiating ART early after infection (transient reservoirs) versus late (persistent reservoirs). Using immunoPET/CT-guided sampling of SIV-infected rhesus macaques combined with spatial transcriptomics, we interrogated tissue sites of viral production during the eclipse phase following analytical treatment interruption (ATI). Our results revealed that viral rebound from persistent reservoirs arises from discrete, transcriptionally active foci enriched in the mucosa lining the gut lumen.

Eclipse phase persistent reservoirs were characterized by increased proviral burden and a distinct tissue state marked by activation of stress-response, metabolic, mitochondrial, and cell cycle programs coupled to repression of cytoplasmic translation and increased cellular senescence. These features co-occurred with immunosuppressive cellular architectures resembling tertiary lymphoid structures enriched for Treg cells, innate lymphoid cells, and mast cells, regulated by Treg-centered cell–cell interaction networks. In contrast, transient reservoirs displayed enhanced translational and metabolic activity and were embedded within immune-active environments enriched for CD8⁺ T cells, Th17, Tfh, and activated CD4⁺ T cells.

Machine learning identified stress adaptation, hypoxia, metabolic rewiring, and cytoskeletal remodeling pathways as dominant predictors of viral density within persistent VMEs, with strong convergence on programs observed in tumor microenvironments (TME). Orthogonal validation confirmed activation of the integrated stress response (ISR) at sites of viral production in concurrence with results of immunofluorescent microscopy revealing SIV gag expression in two populations primarily in the mucosa, differentiated by the phosphorylation of eIF2α. Together, these findings establish the VME as a critical determinant of reservoir persistence, integrating immune regulation, tissue remodeling, and translational control to enable viral survival. This framework suggests that effective HIV cure strategies will require coordinated disruption of VME-supportive functions in addition to targeting infected cells.

## INTRODUCTION

HIV reservoirs are seeded extremely early after infection and persist primarily within mucosal and lymphoid tissues during prolonged, viremia-suppressing antiretroviral therapy (ART)^1–3^. Gut-associated lymphoid tissues (GALT) along the gastrointestinal (GI) tract harbor the largest of these reservoirs, reflective of its relatively high abundance of susceptible target cells^4,5^. However, the cellular and molecular features of the tissue microenvironment that enable HIV persistence in GALT remain incompletely defined, constraining efforts to achieve durable remission or cure.

Characterization of persistent HIV reservoirs in human tissues has been limited by the feasibility of finding these sites and the need for invasive sampling^6^. In contrast, extensive tissue sampling around analytical treatment interruptions (ATI) in SIV-infected non-human primates have offered a powerful window into reservoir biology^7–12^. ATI allows the infection of new cells to proceed, spreading into local tissue during the eclipse phase, before the amplifying virus spills over into the blood to be measured as viremia^13,14^. The increasing replication is highly focal within gut tissues and shaped by local immune pressures before viremia is evident^15,16^. Furthermore, a pivotal study by Okoye et al. has shown that the timing of ART initiation relative to virus acquisition profoundly alters reservoir fate and composition. ART initiation very early after acquisition (*i.e.*, 4 to 5 days or less) results in a smaller “transient” reservoir that can rebound with ATI after 6 months on ART, but is cleared without rebound if instead ATI occurs after one year of ART suppressed viremia^17^. In contrast, a delay in ART initiation until day 6 leads to a majority of animals rebounding with ATI after a year. Initiation of ART on day 8 or later, almost invariably results in the formation of a robust, “persistent” reservoir that reliably drives viral rebound upon ATI. This study reveals that time is an essential component of the development of the reservoir of persistence. Other studies have also suggested the same conclusion^18,19^. Understanding the differences between these transient and persistent viral reservoirs through their definition shortly after ATI (*i.e.*, during the eclipse phase) may reveal novel curative strategies. Unfortunately, direct interrogation of these sites containing tissue reservoirs has remained exceptionally challenging as virally infected cells during ART and the eclipse phase are rare, spatially constrained, and difficult to identify^20^.

To overcome these barriers, we leveraged an immuno-positron emission tomography/computed tomography (immunoPET/CT)-guided approach^21–23^ to identify the earliest sites of viral rebound within the gut of SIVmac239-infected rhesus macaques (RM) shortly after ATI for characterization by spatial transcriptomics and other assays. This timing takes advantage of the rebounding virus after ATI, which increases the number of infected cells proximal to the reservoir site. Amplification of viral proteins expression facilitates our ability to identify the small piece of tissue containing the viral foci. As the ART fades, the virus can infect nearby susceptible cells and through multiple rounds of replication potentially alter the resident tissue erasing or masking the footprint of the reservoir of persistence that provided the spark. By looking at the site of the rebound very early, in the eclipse phase before the reemergence of systemic viremia, it is possible to gain insight into the reservoir structure and composition before ATI. Theoretically, the foci of early rebound should contain two populations of SIV infected cells, those associated with the reservoir of persistence which have survived for more than a year, and the newly infected cells that will be the origin of the population that will be shaped by host responses and local immune pressures before viremia becomes evident. By comparing transient and persistent reservoir sites during the eclipse phase in animals that started ART at different times after SIV infection, we defined spatially resolved cellular programs and tissue microenvironmental features that were common to both or distinguished the two states. Both reservoir conditions were characterized by epithelial cell-associated GI tissue regions and increased transcription specifically associated to SIV presence. Our results indicated that sites of rebound within animals with persistent reservoirs were in cellular environments resembling immunosuppressed tertiary lymphoid structures (TLS)^24,25^ transcriptionally characterized by regulatory T cells (Treg), mast cells, innate lymphoid cells (ILC), and chronic stress response programs with repressed translational capacity –a known consequence of the inhibition of global, cap-dependent translation caused by activation of the integrated stress response (ISR)^26–28^. In contrast, sites of rebound within animals with “transient” reservoirs had more infiltrated adaptive effector cell populations and were characterized by enhanced translational and metabolic programs consistent with activated immune responses^29,30^. These results suggest a model in which untreated SIV/HIV infection promotes the formation of a specific virus microenvironment (VME) within the GALT that enables persistence during therapy though immune exclusion, limited antigen production, and cellular senescence. Like the analogous tumor microenvironment (TME)^31^, the VME would support the survival of the infected cells while limiting hostile host responses seeking their elimination. This new paradigm establishes a framework towards the development of a functional cure for HIV by achieving viral clearance through simultaneous therapeutic disruption of the multiple VME reservoir-supportive functions while targeting the infected cells that compose the persistent reservoir.

## RESULTS

### ImmunoPET/CT-guided spatial transcriptomics allows for specific analysis of SIV reservoirs

Throughout this study, we compared initial sites of viral rebound early after ATI in tissue resections from SIV infected RMs that either received ART early after SIV acquisition (modeling a “transient” reservoir) or late after SIV acquisition (modeling a “persistent” reservoir) (**Supp. Fig. 1A**). Animals in the Early ART / Early ATI group (EAEA, n=4) were intravaginally/intrarectally challenged with SIVmac239 and ART was initiated at 4 days post-infection. ART was continued for 6 months followed by ATI and euthanasia 4-5 days later, prior to detectable viremia (**Supp. Fig. 1B**). In contrast, animals in the Late ART / Early ATI group (LAEA, n=3) were intravenously challenged with SIVmac239 and ART was initiated at 10 weeks post-infection. ART was continued for 12 months followed by ATI and euthanasia 4-6 days later. Two of the LAEA animals had no detectable viremia at the time of euthanasia and one had a low viral load of 1.2 × 10^−3^ copies/mL (**Supp. Fig. 1B**).

The day before euthanasia, each animal was intravenously administered a ^64^Copper-labelled antibody-derived Fab2 probe recognizing the SIV Env protein to enable identification of localized foci of viral antigen production within the tissues via immunoPET/CT (**Supp. Fig. 1C**). After necropsy, tissues were imaged, partitioned into smaller sections containing positive probe signal, and placed into cryomolds for tissue sectioning. To first validate our approach, three axially adjacent (separated by 20µm) tissue sections from the transverse colon from one EAEA animal were processed either for spatial transcriptomics on the 10x Visium platform, for proviral Quantitative Real-Time PCR (qPCR), or for immunofluorescent (IF) staining with antibodies against SIV Gag and immune cell markers (**Supp. Fig. 2A**). The spatial transcriptomics data was subjected to unsupervised clustering (**Supp. Fig. 2B**) and differential gene expression analysis to identify cluster-specific markers. Cluster identities were overlaid onto the Hematoxylin and Eosin (H&E) stained image (**Supp. Fig. 2C**).

Paired IF imaging revealed several foci of viral protein production per section, consistent with prior observations during the eclipse phase (**Supp. Fig. 2A**). The overall morphology of the adjacent tissue sections was highly comparable and was recapitulated by the spatial transcriptomics data. For example, an immune aggregate visible on the H&E stain (purple circle, **Supp. Fig. 2A**) had a high frequency of CD3+ cells and no detectable SIV Gag expressing cells. This immune aggregate coincided with a single transcriptional cluster (cluster 6; **Supp. Fig. 2B-C**) that was highly enriched in genes associated with gene ontology (GO) terms related to immune processes (**Supp. Fig. 2D**). Similarly, cluster 3, which overlapped with areas displaying high density of viral foci in the IF staining (red circle, **Supp. Fig. 2A**), was characterized by differential enrichment for GO terms associated with viral production (**Supp. Fig. 2E**). Taken together, these data validate the use of immunoPET/CT for the identification of tissue blocks containing localized foci of viral antigen expression for downstream characterization by IF imaging and spatial transcriptomics.

### SIV antigen production during the eclipse phase of ATI is associated with higher levels of host transcription

Based on high levels of viral antigen detected by immunoPET-CT in each block, 17 colon and one jejunum tissue sections (11 from 3 LAEA animals, 7 from 4 EAEA animals) were selected for spatial transcriptomics. An axially adjacent tissue section from each block was subjected to IF staining for SIV Gag expression. Overlay of the IF image with the spatial transcriptomics grid enabled us to classify SIV-positive versus SIV-negative spots (**Figure 1A**, center; refer to the **Methods** and **Supp. Fig. 3**). Comparing these different regions showed that overall host transcription levels were significantly higher in SIV-positive than SIV-negative spots in both EAEA and LAEA animals (**Figure 1B**). Among EAEA tissues, SIV-positive spots had a median of 8,633 reads, while SIV-negative spots had a median of 4,318 reads (p-value = 3.78e-03). Among LAEA tissues, SIV-positive spots had a median of 9,827 reads and SIV-negative spots had a median of 5,967reads (p-value = 3.09e-09).

**Figure 1.**
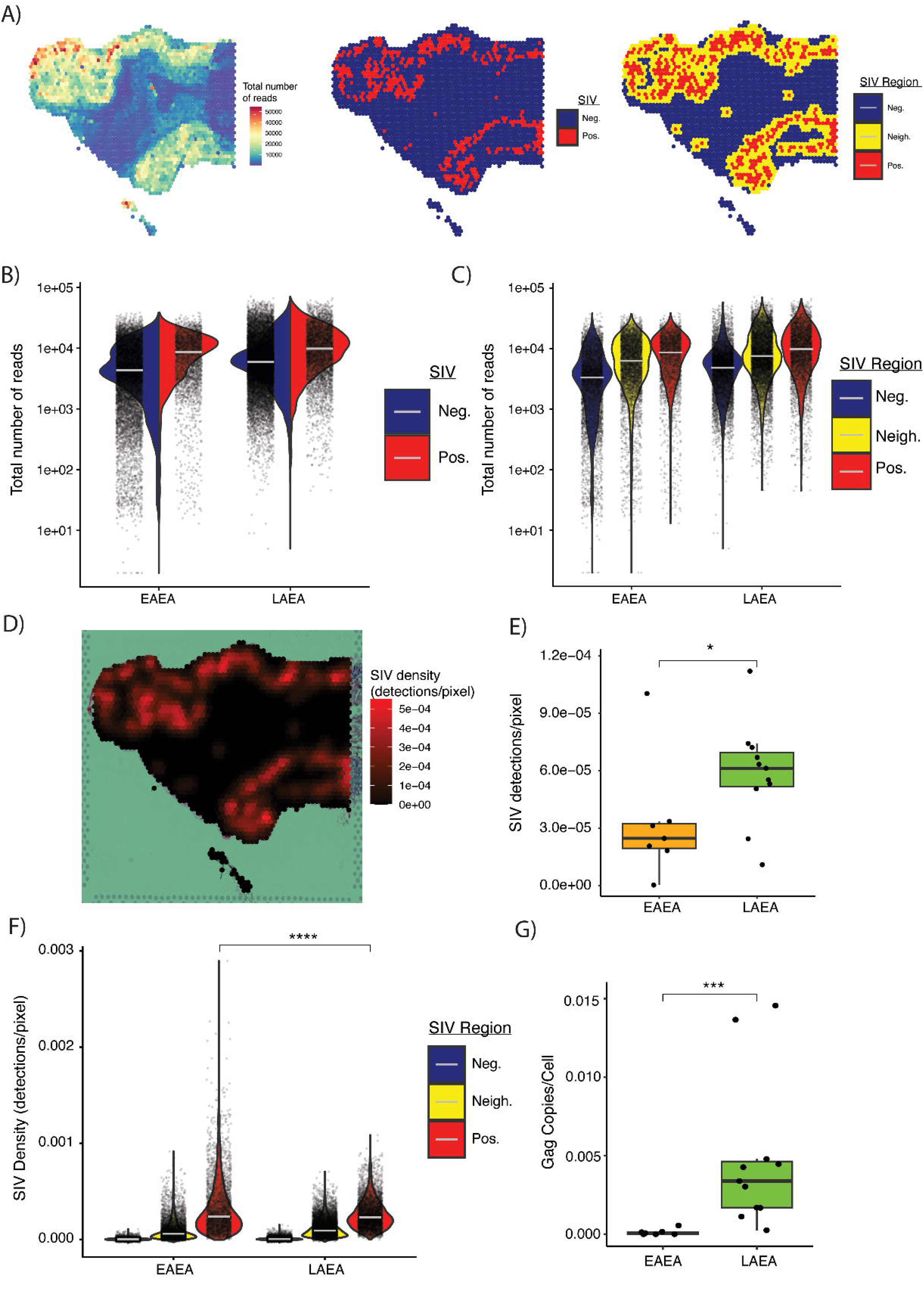
Spatial transcriptomics associated to SIV presence during early rebound. **A)** Spatial distribution of transcriptional reads (left), SIV Negative vs. SIV tags (center), and SIV Negative, SIV Neighbor, and SIV tags (right) at each spatial transcriptomics spot in a slide from the persistent reservoir condition. **B)** Violin plot representing the numerical distribution of reads at each spot, split by SIV Negative and SIV Positive spots for EAEA and LAEA groups. **C)** Violin plot representing the numerical distribution of reads at each spot, stratified by SIV Negative, SIV Neighbor, and SIV conditions for each of the animal groups. **D)** SIV density, obtained by converting SIV presence in spatial transcriptomics spots from a categorical variable to a quantitative measure by fitting IF pixel intensities to a density function, is visualized on a slide from the persistent reservoir condition. **E)** Boxplot representing the median SIV density measured as detections/pixel for each slide in the transient and persistent reservoir conditions (EAEA vs. LAEA). **F)** Violin plot representing the numerical distribution of SIV density associated to each spot, stratified by SIV Negative, SIV Neighbor, and SIV conditions for each of the animal groups. The comparison between SIV density specifically at SIV positive spots (controlling for time since ATI) is indicated by the bracket. **G)** Boxplot representing the proviral levels per animal group as measured by Gag qPCR or dPCR in adjacent tissue sections to the ones analyzed using spatial transcriptomics. For all violin plots dots represent each Visium spot, and grey line represents the median for each condition represented by the violin. For all boxplots each dot represents a slide, the boxes represent the median and interquartile range (IQR), and the whiskers extend from the highest and lowest values no further than 1.5*IQR.

To determine if this increased transcriptional activity was specifically associated with SIV antigen presence or rather a reflection of the tissue architecture in these regions (*e.g.*, mucosal regions vs. muscle rich regions), we added an additional designation for SIV-negative spots that were immediately adjacent to SIV-positive spots (*i.e.*, “SIV-neighbors”). This designates a third region of the tissue surrounding the foci of SIV production (**Figure 1A**, right and **Supp. Fig. 3**). Repeating the same analysis combining both animal groups, we see significantly higher transcriptional levels in SIV-positive spots compared to both SIV-neighbor spots (LAEA p-value = 0.0130; EAEA p-value = 0.0298) as well as SIV-negative spots (LAEA p-value = 6.993e-06; EAEA p-value = 0.0005) (**Figure 1C**). However, SIV-neighbor spots also had significantly higher transcriptional levels than SIV-negative spots (LAEA p-value = 0.0017; EAEA p-value = 0.0202), consistent with the clear morphological patterns in overall read count (**Figure 1A**, left). This suggests that both SIV detection and tissue composition/architecture is associated with increased cellular transcription.

We next analyzed if the distribution of the SIV-positive spots was random or significantly clustered in specific regions of the tissues. Clustering was visually evident (*e.g.*, **Figure 1A**) in all slides in both groups and statistically significant (quadrat test of Complete Spatial Randomness p-value < 0.0001), with no detectable differences in the degree of such clustering between the EAEA and LAEA groups. This result indicates that viral antigen production is localized in specific tissue regions during the eclipse phase of an ATI. This is consistent with viral antigen production early during the eclipse phase of an ATI remaining within specific, spatially constrained foci, rather than initial viral reactivation off-ART being stochastic across all proviral DNA-containing cells in tissues.

### Later ART initiation is associated with more foci of antigen detection, but less antigen production early after ATI

To complement the previous analysis, the SIV Gag signal associated with each Visium spot was quantified using the IF signal density value per pixel of the image coordinate closest to the center of each spot (**Figure 1D** and **Supp. Fig. 3**). The median SIV Gag signal density across all pixels in each slide was significantly higher in LAEA animals compared to EAEA animals (p-value = 0.034) (**Figure 1E**). To determine if this was due to the number of spots or the intensity of the signal within each spot, the median SIV Gag signal density per pixel in SIV-positive, SIV-neighbor, and SIV-negative spots was calculated (**Figure 1F**). In both LAEA and EAEA animals, spots labeled as SIV-positive had the highest SIV Gag density, followed by the SIV-neighbor spots, while SIV-negative spots had very low values (**Figure 1F**). Notably, although there were more SIV-positive spots in the LAEA animals compared to EAEA animals (LAEA total SIV-positive = 5746 vs. EAEA total SIV-positive = 3725), the SIV Gag signal density in the SIV-positive spots was significantly lower (p-value = 1.97e-13; **Figure 1F**). This reveals that there are more SIV Gag antigen foci in the LAEA animals compared to the EAEA animals, but that each foci produces less antigen.

To determine if there are more cells harboring SIV proviruses in the LAEA compared to EAEA tissue sections, SIV proviral DNA copies were quantified in the slice of tissue immediately adjacent to those analyzed by spatial transcriptomics. Indeed, qPCR for integrated SIV Gag copies showed higher copies in LAEA tissues compared to EAEA tissues (LAEA mean: 1.27e-03 gag DNA copies/cell; EAEA mean: 4.81e-04 gag DNA copies/cell; Wilcoxon p-value= 0.0008; **Figure 1G**). These results indicate that animals with a persistent reservoir have more integrated proviruses and see more sites of SIV antigen production during the eclipse phase of ATI, but produce less antigen per foci relative to animals with a transient reservoir.

### Differential host gene expression pathways across SIV-positive, -neighbor and -negative gut tissue spots during early ATI after either late or early ART initiation

To characterize the transcriptional signatures associated with early ATI, we examined differentially expressed genes (DEGs) and significantly enriched pathways across the three spot categories (methodological details in **Figure 2** legend), starting with LAEA tissue SIV-positive spots. Pathways associated with HIV infection, HIV lifecycle, and host interactions with HIV factors were upregulated in LAEA tissue SIV-positive spots (**Supp. Fig. 4 and Supp. Table 1**). This independently confirmed the specificity of spots identified as SIV-positive by integration with IF SIV Gag detection. Other upregulated transcripts in LAEA tissue SIV-positive spots included DEGs related to cell cycle and cell proliferation –e.g., E2F and MYC targets, DNA synthesis and replication (G2/M checkpoint), metabolic activation, RNA processing, and mitochondrial translation. Strong activation of DEGs related to cellular stress responses were also prominent, such as the unfolded protein response to endoplasmic reticulum (ER) stress and upregulation of GCN2 in response to amino acid deficiency. In LAEA SIV-positive spots, strong down-regulation –i.e., activation in SIV-negative regions–of extracellular matrix (ECM) pathways was also found, including ECM organization, collagen biosynthesis and formation, integrin interactions, and focal adhesion.

**Figure 2.**
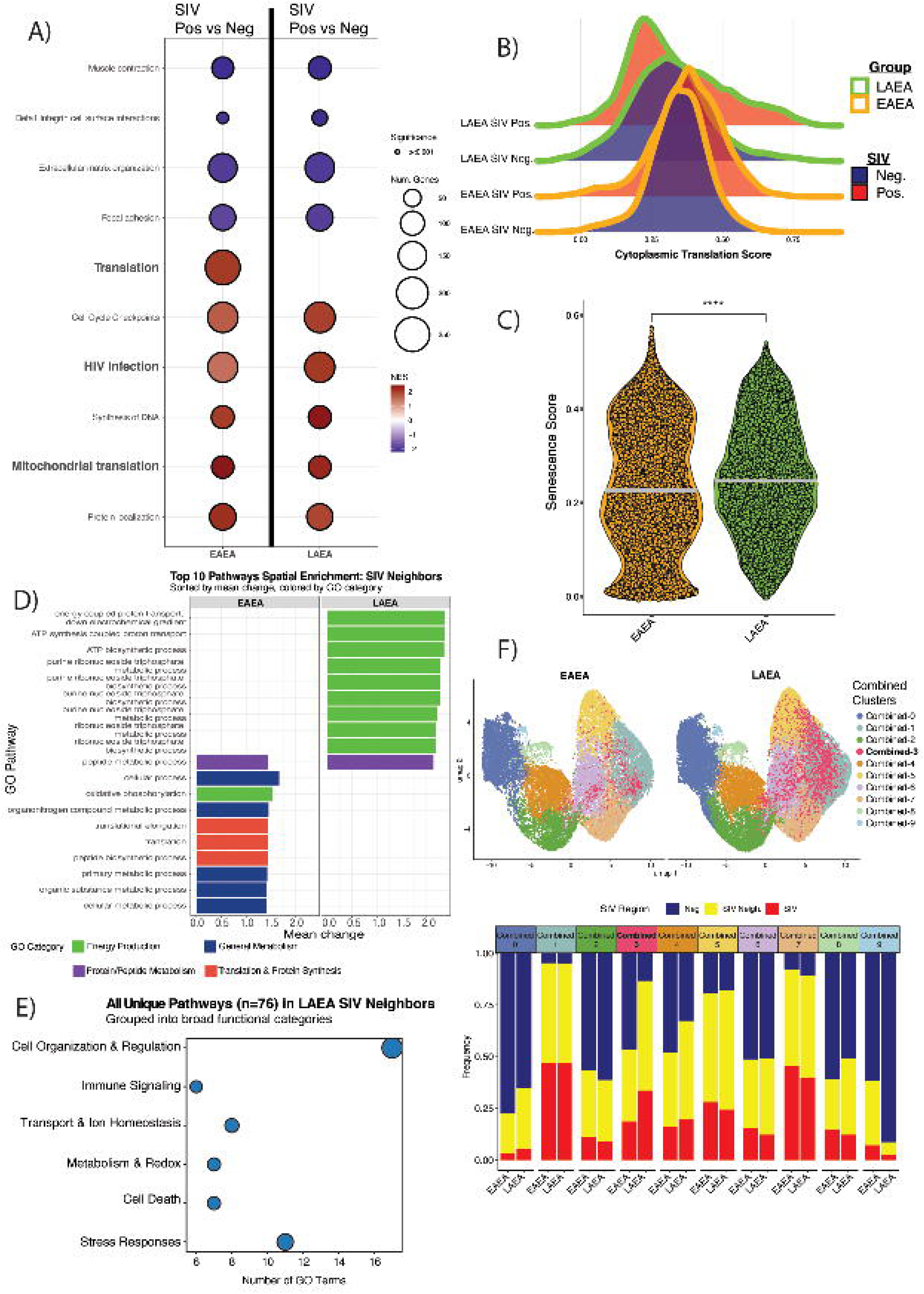
Differential gene expression and spatial analysis of transcriptional patterns in transient vs. persistent reservoirs. **A**) GSEA analysis among canonical pathways and hallmark gene sets from MSigDB and GO, comparing EAEA and LAEA. After MAST analysis of differential expression across the 3 spot categories (controlling for the slide of origin of each spot), GSEA was performed using the canonical pathways subcollection and the Hallmark gene sets included in the macaque MSigDB and the macaque GO databases; analyses used the entire list of transcripts that appeared in at least 1% of the spots and they were ranked by the log-fold change between SIV-positive and SIV-negative spots. **B**) Distributions of cytoplasmic translation scores, grouped by transient vs. persistent reservoir conditions and SIV Negative vs. SIV spots. **C**) Senescence score distribution among all spots in the transient vs. persistent reservoir conditions. The persistent reservoir was associated with higher senescence scores, a marker of cell stress responses. **D**) Top 10 pathways attained by spatial enrichment analysis of GO Biological Process pathways comparing SIV neighbor spots in LAEA and EAEA. **E**) Distribution of broad GO functional categories among all pathways that were uniquely enriched in LAEA SIV-neighbor spots. **F**) UMAP visualization of transcriptional clusters in spatial transcriptomics data and distribution of SIV presence in each transcriptional cluster. Cluster detection was consistent among the integrated data for the transient and persistent reservoir conditions, but the distributions of the presence of SIV within clusters were more heterogeneous in the persistent reservoir condition than the transient reservoir condition.

Similar differential expression patterns were observed in EAEA tissue SIV-positive spots, including both upregulation of cell cycle and proliferation gene expression as well as downregulation of ECM related pathways in comparison to SIV-negative spots. HIV infection-related transcripts were also upregulated, albeit less so than in LAEA SIV-positive spots; this is consistent with lower provirus DNA levels detected adjacent to EAEA than LAEA SIV-positive spots. Some transcriptional pathways significantly differed between LAEA and EAEA SIV-positive spots. Most notably, EAEA SIV-positive spots displayed high levels of overall translation activation as well as mitochondrial translation activation; only the latter was observed in LAEA SIV-positive spots (**Figure 2A**). This difference in translation activation was confirmed by comparing a cytoplasmic translation score among the 4 categories of EAEA and LAEA spots with either SIV positivity or negativity (**Figure 2B**); LAEA SIV-positive spots had the lowest levels of cytoplasmic translation. Because cell cycle arrest and stress responses were also more notable in LAEA SIV-positive spots, cell senescence scores were determined using the senePy database and analysis pipeline^32^ and compared between all Visium spots in LAEA versus EAEA tissues. This analysis showed higher senescence scores in LAEA tissues (**Figure 2C**; LAEA mean = 0.253; EAEA mean = 0.234; Wilcoxon p-value < 2.2e-16), as well as a different distribution of such scores in LAEA versus EAEA tissues (**Figure 2C**; LAEA IQR = 0.183; EAEA IQR = 0.222).

SIV-negative spots were also studied in both sets of tissues. In both LAEA and EAEA tissue SIV-negative spots, there was upregulation of ECM related pathways, including cell adhesion and structure, Epithelial-mesenchymal transition (EMT), and differentiation programs for myogenesis and angiogenesis; these pathways were each upregulated, in contrast to their downregulation in SIV-positive spots, in both LAEA and EAEA tissues.

The above differences in DEGs pathway enrichment in EAEA versus LAEA SIV-positive and SIV-negative spots inspired an analysis that included the neighboring spots forming a boundary between each spot category in LAEA versus EAEA tissues. This approach maximized the use of spatially resolved transcriptional data through spatial enrichment analysis as described in Methods and **Figure 2D** legend and resulted in the identification of pathways significantly enriched for each tissue region per group (FDR adjusted p-value< 0.05 and absolute mean change> 0.5). LAEA tissues displayed a stronger activation of different pathways in SIV-positive and -neighbor spots compared to EAEA tissues, with the largest differences in these LAEA-unique pathways detected in SIV-neighbor spots (**Figure 2D**). SIV-neighbor spots in LAEA tissues, relative to SIV-neighbor spots in EAEA tissues, had higher levels of energy production and secretory processes. Importantly, multiple stress responses, such as general response to stress, to chemical stress, to amino acid starvation, to oxidative stress, or to unfolded protein (**Figure 2E and Supp. Figure 5**) were also uniquely enriched in LAEA tissue SIV-neighbor spots. In contrast, SIV-neighbor spots in EAEA showed a significantly stronger activation of genes related to translation-associated processes (red bars in **Figure 2D**) and metabolic processes than did those in LAEA tissue SIV-neighbor spots. Some pathways were also uniquely activated in SIV-negative spots in EAEA tissues and not in SIV-negative spots in LAEA tissues (137 unique processes in EAEA SIV-negative spots vs. 46 in LAEA SIV-negative spots; **Supp. Table 2**). EAEA tissues had a more balanced activation of processes amongst the 3 different spots (SIV-positive, SIV-neighbors and SIV-negative).

In summary, key differences in the transcriptional profiles of the reservoir microenvironment between persistent and transient tissue reservoirs during the eclipse phase of an ATI preceding viremia included higher levels of cellular senescence, greater activation of stress responses, extensive activation of biosynthetic processes, and significantly lower levels of cytoplasmic translation in both SIV-positive and -neighbor spots in LAEA versus each of those two corresponding categories of spots in EAEA tissues.

### Different transcriptional clusters are associated with LAEA versus EAEA SIV-positive spots

After observing some differences, as well as many similarities, between transcriptional pathways in SIV-positive and -neighbor spots in LAEA versus EAEA tissues, an in-depth comparison of the specific transcriptional clusters associated with SIV-positive and -neighbor spots was also performed. These transcriptional clusters will be described; first for LAEA tissues and, secondly, for EAEA tissues. Following that, clusters defined in an integrated analysis of the combination of LAEA and EAEA tissues will be examined.

In LAEA tissues, 2 out of 8 transcriptional clusters included the highest levels of SIV-positive spots amongst all their clustered spots compared to the other LAEA clusters (**Supp. Fig. 6**). These LAEA tissue-specific transcriptional clusters LAEA-1 and LAEA-6, included 40.1% and 39.2%, respectively, SIV-positive spots among all their assigned spots (**Supp. Fig. 6E**). Additionally, these clusters contained a very limited number of SIV-negative spots (9.2% and 14.0% respectively), consistent with their enrichment in SIV-positive spots. Stress responses and HIV infection-related transcriptional pathways were upregulated in both these clusters. These two different SIV-associated clusters also each displayed distinct activation of different pathways and biological processes, suggesting that SIV-associated regions in LAEA tissues were heterogeneous in their transcriptional patterns. Relative to cluster LAEA-6, cluster LAEA-1 included significantly higher activation of DEGs associated with the cell cycle, metabolism, and morphogenesis, as well as a stronger association with host factors that interact with HIV-1. In contrast, immune activation signals, such as overall immune responses, cytokine signaling, and activation of both the innate and adaptive immune responses, stood out more prominently in cluster LAEA-6.

Interestingly, transcriptional cluster LAEA-2 was mainly comprised of SIV-neighbor spots (55.7%). When compared to all other clusters, LAEA-2 had a unique set of activated DEGs related to cell-cell interactions and specific activation of innate immune system-related gene expression, indicating distinct characteristics of the regions surrounding SIV foci. On the other hand, other clusters had a much lower frequency of SIV-positive spots, especially cluster 1 with less than 5% of SIV spots.

Among EAEA tissues, clusters EAEA-5 and EAEA-6 contained 51.6% and 44.5%, respectively (**Supp. Fig. 6F**), SIV-positive spots. In comparison, for cluster EAEA-0, only 3.1% of its assigned spots were SIV-positive. In general, there was less heterogeneity, between EAEA SIV-associated clusters (EAEA-5 and EAEA-6) than seen among the LAEA SIV-associated clusters. However, some differences were also noted between these two clusters. Cluster EAEA-5 showed higher upregulation of cell cycle and metabolic pathways, while cluster EAEA-6 had increased signals of morphogenesis related pathways. This was indicative of less heterogeneity within the SIV-associated clusters in transient reservoirs.

Due to the observed differences in transcriptional clusters associated with SIV-positive and -neighbor spots in separate analyses above of persistent and transient tissues, an integrated analysis was performed combining and re-clustering the transcriptional data from the two groups of tissues. This direct comparison confirmed important differences between LAEA and EAEA. While two clusters highly associated with SIV presence were consistently detected in both groups, we detected an unbalanced presence of one of these SIV-associated clusters (cluster Combined-3 in the integrated analysis **Figure 2F** and **Supp. Fig. 7A**) that was significantly more abundant in LAEA where it also displayed higher frequency of SIV-positive and SIV-neighbor spots. This cluster was overall characterized by genes related to GTPase regulator activity and phospholipid metabolism, as well as immune and HIV-related genes. Differential expression analysis between the spots assigned to this cluster from LAEA or EAEA indicated, in concordance with previous results, that downregulation of translation related pathways in LAEA was again the main driver of the observed differences (**Supp. Fig. 7**).

Overall, this analysis indicated a high degree of heterogeneity at the eclipse phase of an ATI preceding viremia in the areas of SIV positivity in persistent reservoirs in LAEA tissues that was not as evident in transient reservoirs in EAEA tissues. SIV-associated clusters in LAEA tissues showed expression of stress response genes, HIV-1 infection-related genes, adaptive and innate immune system activating genes, and downregulation of translation-related pathways. An abundance of transcripts reflecting cell-cell interactions and activation of innate immune system was seen only in LAEA SIV-neighbor spots.

### Associations of inferred broad and immune-specific cell type proportions inform about eclipse phase gut reservoir features and their changes with longer untreated infection

Associations of the inferred proportions of broad Pan-GI cell types from the human gut cell atlas were analyzed across the range of observed SIV antigen signal density in all Visium spots (as described in **Methods**). A strong positive association of inferred proportions of epithelial cells with SIV Gag antigen IF signal density was seen with in both EAEA and LAEA tissues (**Figure 3A, left panel**); epithelial cells are known to be the predominant cell types close to the gut lumen^33^. In contrast, a negative association was observed between proportions of mesenchymal cells with SIV antigen density in both EAEA and LAEA tissues (**Figure 3A, left panel**); mesenchymal cells comprise a larger fraction of the cell types more distal from the gut lumen^33^. The inferred frequency of red blood cells (RBC), and to a lesser extent that of neuronal cells, was negatively associated with SIV antigen density in EAEA tissues and positively associated in LAEA tissues (**Figure 3A, left panel and Supp. Fig. 8**). Concordant directionalities of associations of these 4 non-immune cell types in EAEA and LAEA tissues were seen in different analyses within each of the 3 categories of spots (SIV-negative, SIV-neighbor, and SIV-positive) (**Figure 3A, right panel**).

**Figure 3.**
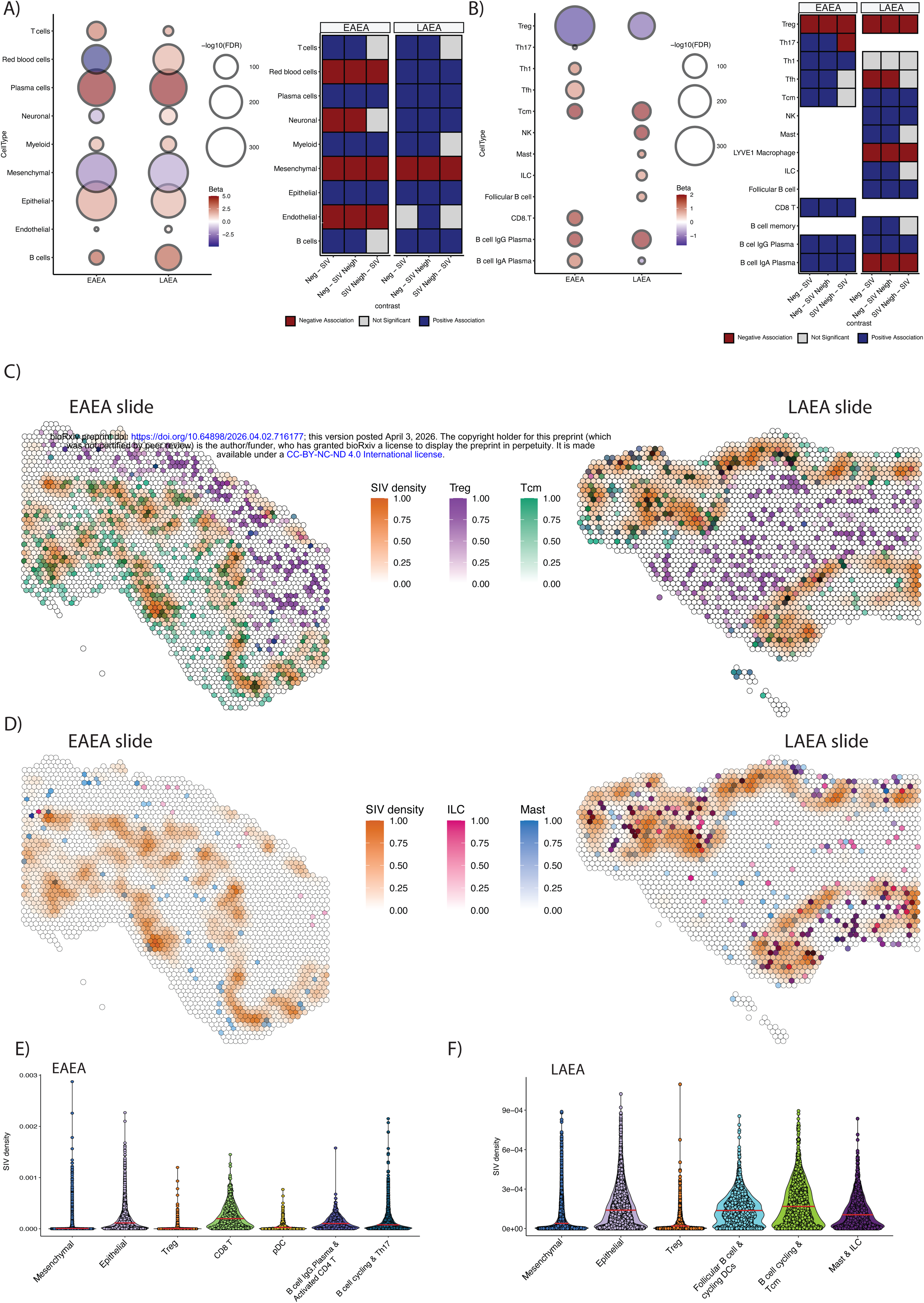
Cell types at each spatial transcriptomics spot inferred via machine learning-based gene expression deconvolution. **A)** Dotplot of inferred cell types for broad classes of gut cells (Gut Cell Atlas) in the transient and persistent reservoir conditions, indicating cell type associations with the presence of SIV foci of infection. Heatmap of pairwise statistical comparisons of frequencies of inferred cell types in Negative, SIV Neighbor, and SIV conditions. **B)** Dotplot of inferred cell types for specific sub-classes of gut immune cells (Colon Immune Atlas) in the transient and persistent reservoir conditions, indicating cell type associations with the presence of SIV foci of infection. Heatmap of pairwise statistical comparisons of frequencies of inferred immune cell sub-types in Negative, SIV Neighbor, and SIV conditions. **C)** Cell frequency and SIV density spatial overlay on example slides from the transient and persistent reservoir conditions, visualizing a negative association between SIV density and regulatory T cells (Treg) and a positive association between SIV density and central memory T cells (Tcm). **D)** Cell frequency and SIV density spatial overlay on example slides from the transient and persistent reservoir conditions, visualizing the positive association between SIV density and mast and innate lymphoid cells (ILC) in the persistent reservoir that was not present in the transient reservoir condition. **E)** Associations of spatial distribution of the proximity of cell types to SIV, as quantified by SIV density, in the transient reservoir condition. **F)** Associations of spatial distribution of the proximity of cell types to SIV, as quantified by SIV density, in the persistent reservoir condition.

Subsequently, we focused on inferring immune-specific cell types using the human colon immune cell atlas as background (as noted in **Methods**). Results of these immune cell analyses will be detailed here and then features of eclipse phase gut reservoirs they suggest will be described. Overall, the two animal groups showed a strong significant negative correlation between Treg and SIV density (**Figure 3B, left panel)**; this negative correlation of Treg cells with SIV positivity was also seen in separate analyses of each of the spot categories in either LAEA or EAEA tissues (**Figure 3B, right panel; Figure 3C**). Both groups also showed a significant positive correlation between Tcm and SIV density (**Figure 3B, left panel; Figure 3C**), with LAEA showing a more defined Tcm gradient when comparing the spot categories (**Figure 3B, right panel**). Also, IgG-producing plasma B-cells were associated with SIV positivity in both EAEA and LAEA tissues (**Figure 3B, both panels**). Instead, cells such as mast cells, ILCs, natural killer cells (NK), and follicular B-cells were each positively associated with SIV antigen density only in LAEA tissues, while none of these cell types were found to be associated or consistently detected in the EAEA samples (**Figure 3B, left panel**). Tissue sections showed increased proportions of both mast cells and ILCs in and nearby high SIV density regions in LAEA tissues; these tissues also showed closer proximity of these 2 innate immunity cell types to each other (including some overlap of both cell types in the same spot) and to spots with the highest SIV antigen density (**Figure 3D**). On the other hand, key T-cell subsets were not consistently detected in LAEA while they showed significant associations with SIV levels in EAEA. Among these, CD8+ T-cells and CD4+ Th1 cells were positively and significantly associated with SIV density exclusively in EAEA tissues in the analysis across all spot categories (**Figure 3B, left panel)** and also in separate analyses of each of the spot categories (**Figure 3B, right panel**).

Hypotheses tested in experiments that will be described in following sections were raised by associations between inferred proportions of non-immune and immune cells with SIV Gag antigen density including: that SIV antigen production during the eclipse phase of an ATI is preferentially localized close to the gut lumen; that SIV antigen-producing cells become more proximal to TLS during a longer time left untreated; that adaptive immune responses to SIV are less in LAEA than EAEA tissues; and that a longer duration of untreated infection fosters a more immunosuppressive viral microenvironment.

### Principal components analyses of inferred cell types and transcriptional clusters extend characterization of LAEA versus EAEA eclipse phase tissue reservoirs

Principal components analysis (PCA) further explored the clusters of inferred cell composition and host transcription in microenvironments of EAEA and LAEA tissues. PCA identified components contributing to clusters attained for the Pan-GI broad cell types or the human colon immune cells, as well as to associations of the inferred cell type clusters with distinct transcriptional clusters in Visium spots in each tissue type (**Supp. Fig. 9** details all cell types and associations). The latter analyses added the examination of features that differed in early ATI between EAEA versus the LAEA reservoir of persistence.

With PCA from LAEA tissues we observed that the main contributors for the different clusters observed using the broad gut cell types were differences in frequency of mesenchymal cells versus frequency of epithelial and RBC (**Supp. Fig. 10** where epithelial and RBC are drivers for the broad cell PCA-cluster 3, shown in purple), confirming results shown in **Figure 3**. Regarding the human colon immune cells inferred, each of the clusters were mainly determined by few cell types as indicated in **Supp. Fig. 11D**.

Combining these results with the LAEA-specific clusters previously attained, we could determine the inferred cellular composition of each of the transcriptional clusters. The two transcriptional clusters associated with SIV antigen positivity (LAEA-1 and LAEA-6) each had different inferred cellular compositions (**Supp. Fig. 11E** and **F**). LAEA-1 spots were mostly associated with areas inferred to be characterized either by follicular B-cells, cycling dendritic cells (DC), NK, and memory B-cells (43%) or central memory T-cells (Tcm), cycling B-cells, and IgG B-cells (37%) (**Supp. Fig. 11E**). LAEA-6, with activated immune response transcripts, was more prevalent in areas characterized by Tcm, cycling B-cells, and IgG B-cells (61%) (**Supp. Fig. 11E**), which was comprised of cell types known to be enriched in TLS. Interestingly, LAEA-2 – with a unique set transcripts related to cell-cell interactions and innate immune system-related genes that was at high frequency in SIV-neighbor spots – was mainly associated with areas rich in mast cells and ILCs (**Supp. Fig. 11E**). LAEA-0, characterized by enrichment of ECM pathways, cell-cell interactions, EMT, and cell adhesion that predominated in SIV-negative spots in LAEA tissues, was almost exclusively associated with the Treg (**Supp. Fig. 11E**).

In EAEA tissues, the inferred cell compositions associated with EAEA-specific transcriptional clusters in SIV-positive spots clearly differed from that of the LAEA tissue SIV-positive spots (**Supp. Figure 11C**). Transcriptional clusters EAEA-4 and EAEA-5 that were enriched in EAEA SIV-positive spots were highly associated with an immune cell composition dominated either by T follicular helper cells (Tfh) presence (EAEA-4) or IgG B-cells and activated CD4 T-cells (EAEA-5). Additionally, the EAEA-0 –with almost no presence of SIV–was associated to Treg, mirroring LAEA-0 in the LAEA tissues.

We performed a third PCA using both broad and immune cells and compared cell composition in LAEA versus EAEA tissues (**Supp. Fig. 12A** and **C**). This analysis confirmed the different components of the LAEA SIV-positive foci were mainly composed of follicular B-cells and cycling DCs; cycling B-cells and Tcm cells; and epithelial cells (**Figure 3F** and **Supp. Fig. 12D**). In LAEA tissue SIV-positive spots, areas defined by higher frequencies of mast cells and ILCs were also seen less frequently (**Figure 3F** and **Supp. Fig. 12D**). In the case of the EAEA tissues, areas enriched with CD8 T-cells and, to a lesser extent, areas enriched in B cell IgG plasma cells and activated CD4 T-cells; epithelial cells; or cycling B-cells and T-helper 17 (Th17) cells, were highly associated with the presence of SIV antigen (**Figure 3E** and **Supp. Fig. 12B**).

Overall, these results suggest that the SIV reservoirs in early, pre-viremia ATI are expressing antigen in epithelial cell-rich luminal regions of the colon and display a heterogeneous and diverse cellular composition, including well-characterized cellular reservoirs like Tcm cells. More importantly, some cell types inferred to be specifically associated with the persistent reservoirs in LAEA tissues closely resemble those in germinal centers within GALT, including TLS that generally are close to gut lumen.

### Markedly different cell-cell interactions orchestrate the immune response early during ATI in LAEA (persistent) or EAEA (transient) tissues

Cell-cell interactions (CCIs) analyses were next evaluated among immune cell types in LAEA versus EAEA reservoirs (as described in **Figure 4** legend). Once CCIs were inferred, interaction patterns and main components were estimated for the two groups. For LAEA tissues, the most frequent interactions occurred between a source (sender) cell located in SIV-positive spots and a receiver cell located in SIV-neighbor spots (16.4%), between pairs of cells located in SIV-neighbors (16.0%), or between cells within SIV-negative areas (14.9%) (**Figure 4A**). Amongst these interactions, the most common source cell type was Treg (9%), followed by ILC (8.3%), while the most frequent receivers were also Treg (11.2%), followed by mast cells (8.5%) (**Figure 4B** and **Supp. Fig. 13**). Finally, the most common interaction inferred was between Treg cells across two SIV-negative areas (**Supp. Fig. 13** and **Figure 4C**). In LAEA tissues, CCIs were also more common between SIV-neighbor spots and less common between two SIV-positive spots or an SIV-positive spot with any other category of spot in either direction (**Figure 4A**). On the other hand, analysis of EAEA tissues indicated very different interaction patterns where the most frequent interactions occurred between either SIV-negative (22.7%) or SIV-neighbor (19.9%) spots (**Figure 4A**). The most frequent source cells in EAEA tissues were Tfh (10.6%), Treg (9.2%), and Th17 (8.5%), while the most frequent receivers were also Tfh (11.4%) and Treg (10.8%) (**Figure 4B** and **Supp. Fig. 13**).

**Figure 4.**
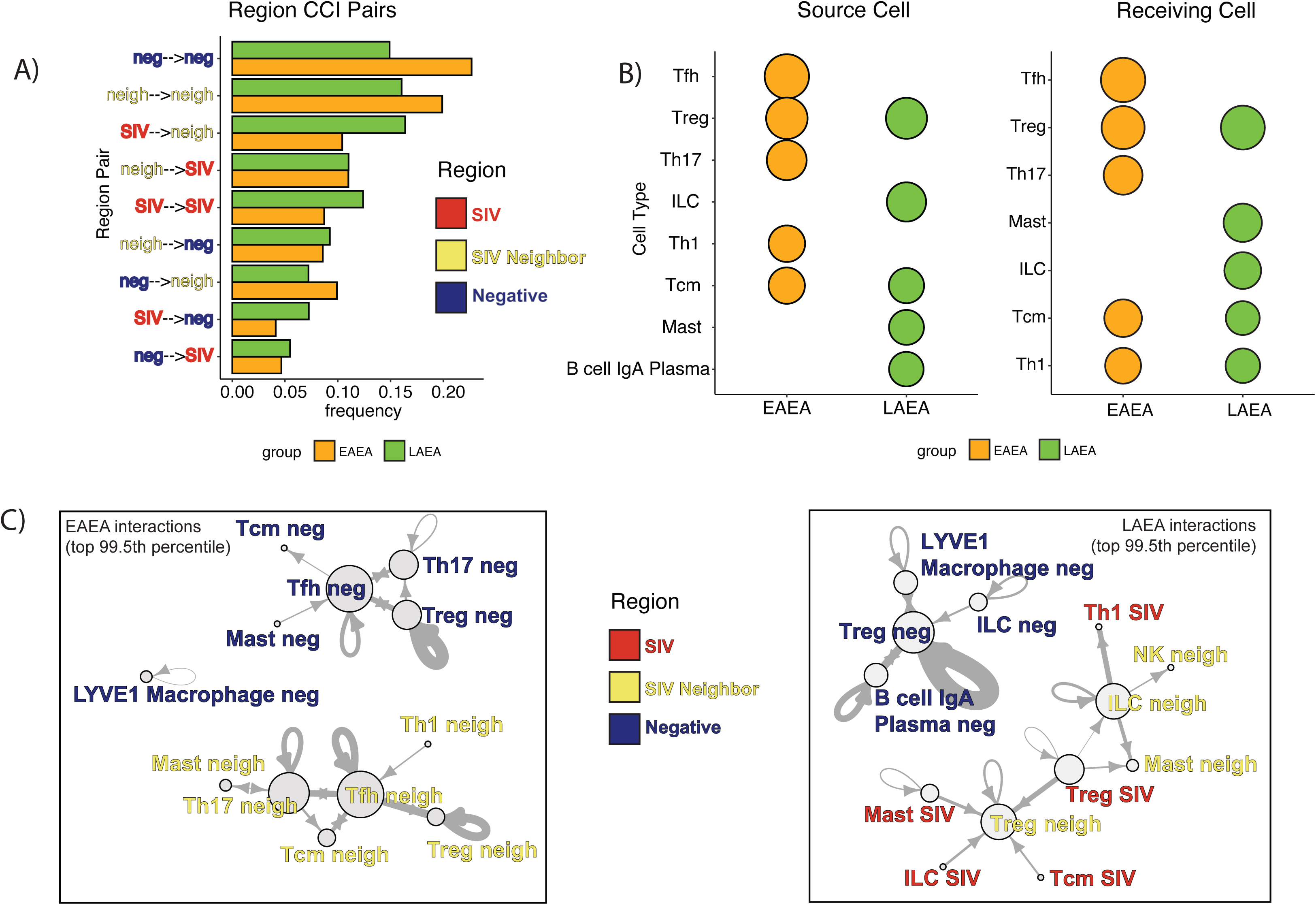
Cell-cell interactions that mediate the immune microenvironment of transient and persistent reservoirs. **A)** Barplots represent the pairwise frequencies at which Negative, SIV Neighbor, and SIV spots interacted within the immune microenvironment for the transient and persistent reservoir conditions. **B)** Dotplot visualizations of the frequency with which immune cell subtypes were the source cell (“sender”) and receiving cell for the transient and persistent reservoir conditions. **C)** Cell-cell interaction network of the top 99.5th percentile of interactions in the transient and persistent conditions, with combined tagging of labels of the interacting cell types with Negative, SIV Neighbor, and SIV labels. Network analyses compared, on a more granular scale, differences in immune responses to rebound initiation in the transient and persistent reservoirs.

Subsequently, an in-depth analysis of the most frequent CCIs inferred for eclipse phase LAEA tissues confirmed the central role of Treg cells in orchestrating the responses to the early rebounding virus from persistent reservoirs (**Figure 4C**). Interactions were very frequent between SIV-positive (**red font in Figure 4A** and **4C**) and SIV-neighbor (**yellow font in Figure 4B** and **4C**) spots or within SIV-negative regions (**blue font in Figure 4A** and **4C**). Interactions between SIV-negative spots and spots in either of the other 2 categories were less frequent (**Figure 4C, right panel**, as indicated by the separately interacting group of only among SIV-negative spot cells in upper left of that panel). In LAEA tissues, Treg cells in SIV-positive spots signaled to Treg cells in SIV-neighbor spots. Additionally, mast cells, ILCs, and Tcm cells in SIV-positive spots were predicted to receive signals from Treg cells in SIV-neighbor spots (**Figure 4C, right panel**). Within SIV-neighbor areas, ILCs were also inferred to play a central role. After being signaled by Treg cells from SIV-positive regions, ILCs can activate other ILCs, mast cells, NK cells, and T-helper 1 (Th1) cells within the same SIV-neighbor region (**Figure 4C, right panel**). Finally, amongst LAEA SIV-negative spots, Treg cells were the most interconnected cell type with possible activation by ILC as well as other Treg (as indicated to arrow looping back to itself in **Figure 4C, right panel**), LYVE-1 macrophages, and IgA plasma B-cells (**Figure 4C, right panel**).

In contrast, the most frequent CCIs in EAEA tissues highlighted Tfh cells as the most centrally located cells in the CCI networks (**Figure 4C, left panel)**. Tfh cells were highly interconnected independently, either within SIV-negative or SIV-neighbor spots in EAEA tissues (**Figure 4C, left panel)**. In EAEA tissues, no interactions were observed between different regions amongst the most common interactions and no CCIs involved cells in SIV-positive areas (**Figure 4C, left panel)**. Inside SIV-neighboring spots, crosstalk within and between Tfh and Th17 cells were the most interconnected CCIs (bottom group of interactions in **Figure 4C, left panel)**. EAEA SIV-neighbor spots also had CCIs between the following: Th1 and Tfh cells, Tfh and Treg cells, and Th17 and Tcm cells (**Figure 4C, left panel)**. Finally, EAEA SIV-neighbor Th17 cells were also found to interact with mast cells and Tcm cells (**Figure 4C, left panel)**. Overall, these results indicate major differences in crosstalk between immune cell types during early ATI in persistent versus transient reservoirs.

### Machine learning modeling identifies the main features of persistent tissue reservoirs

Machine learning approaches were used to identify the top drivers of SIV presence and SIV production levels within the tissue reservoirs. Regression models determined the gene expression most highly associated with SIV levels within tissues from either LAEA or EAEA tissues, followed by classification models to identify the main features that specifically defined SIV-positive areas in LAEA tissues **(Figure 5A and B**). After multiple comparisons, the best performing method was ^25^both for regression and classification models^34^. The regression model identified the transcripts most highly associated with SIV antigen density either for LAEA or EAEA tissues. The main drivers of SIV antigen for LAEA (persistent) reservoirs were significantly different from those for EAEA (transient) reservoirs (**Figure 5A**). For LAEA reservoirs, the 5 most important features in these models were ACADS, ENSMMUG00000018740 (BSG; **Figure 5C**), PFKL, TPM3, and SELENBP1. For EAEA reservoirs, the top 5 features were PIGR, MT1E, RACK1, RPL27A, and ENSMMUG00000059742. However, many of the top features in LAEA reservoirs, appeared in the EAEA reservoir model with lesser importance. Therefore, the genes exclusively found to be expressed in the LAEA model were selected from the top 100 most correlated transcripts. These genes were mainly involved in metabolic regulation, mitochondrial function, immune regulation and membrane trafficking (**Figure 5D**). Interestingly, several of these genes are documented to be highly relevant in cancer, including ERBB2, ERBB3, MAPK6, SGK1, and HOXB13. Cell signaling was also highly represented (PPP4C, PPP2CB, PPM1G, PTP4A2, ETNK1, RIOK3, ATP13A2, STAP2). Additionally, multiple highly correlated genes are involved in mitochondrial function, such as MRPL27, MRPL14, MRPL28, MRPL51, ACAA1, ADCK5, CHCHD1, HSPA9, and HK1. Lastly, metabolic regulators (HMGCR, SCAP, HNF4A, SIRT6, NAPRT, HK1, ACAA1, PNPLA2) and immune regulators (TNFRSF14, VSIR, ZBTB7B, NR2F6, F11R, STAP2) were also found to be important features of the regression model.

**Figure 5.**
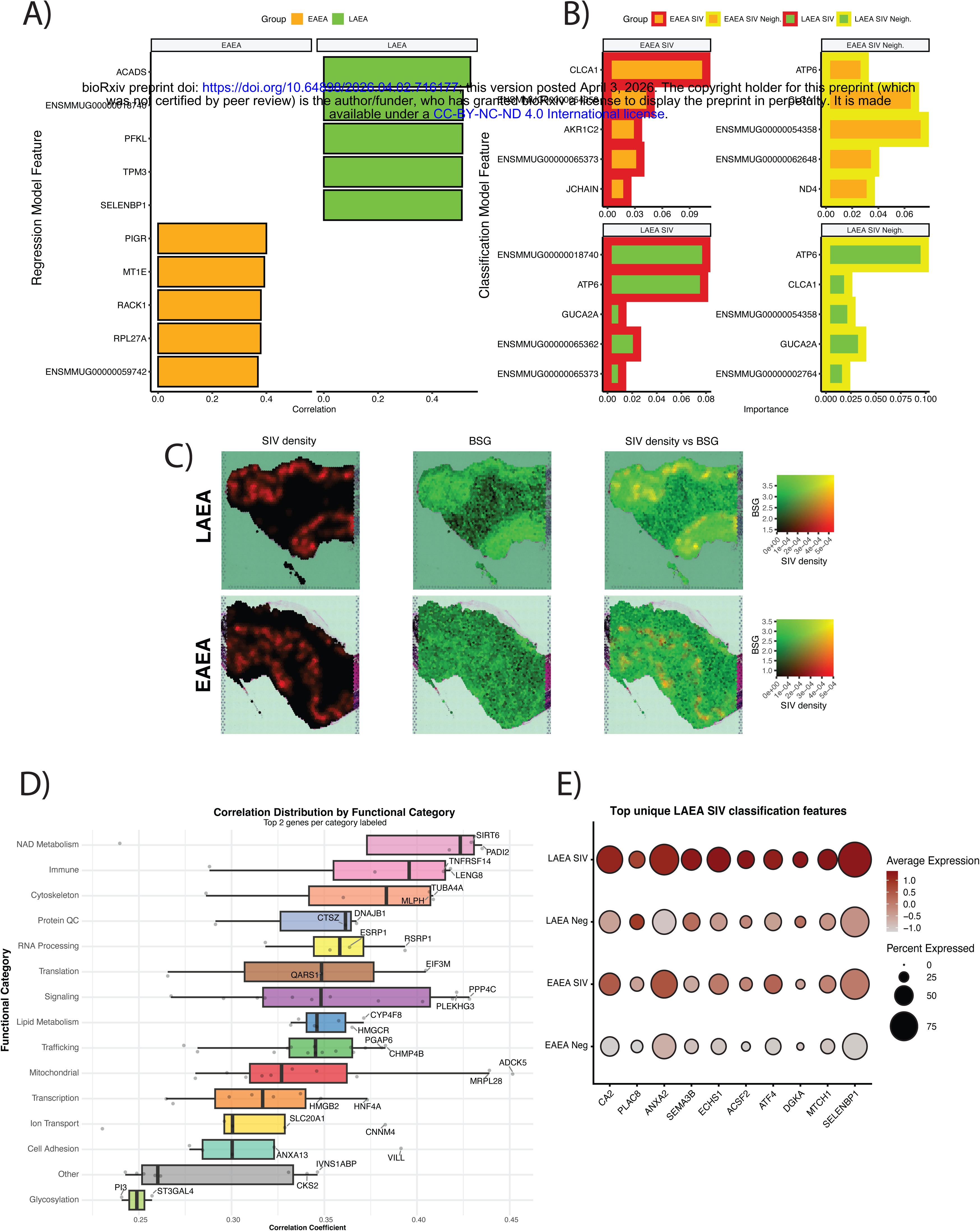
Key gene drivers of viral persistence identified via machine learning. **A)** Regression analysis via a Light Gradient-Boosting Machine Learning model reveals key gene drivers correlated with transient and persistent reservoirs. **B)** Classification analysis comparing SIV vs. SIV Negative or SIV Neighbor vs. SIV Negative conditions via a Light Gradient-Boosting Machine Learning model reveals key gene drivers of importance in transient and persistent reservoirs. **C)** Visualization on a slide from the persistent reservoir condition of the distribution of BSG (ENSMMUG00000018740), the top ranked hit by importance in the persistent reservoir from the machine learning classification analysis, as well as SIV density, both separately and overlaid. **D)** Functional categories of the top 100 key gene drivers exclusive to the persistent reservoir condition, with the correlation coefficient distribution for each category and the top two genes per category labeled. **E)** Dotplot of the top ten gene drivers among the top 75th percentile importance score or above that were unique to the persistent reservoir SIV foci of infection, grouped by category: persistent and SIV, persistent and SIV Negative, transient and SIV, and transient and SIV negative. Average expression and percentage of spots expressing the genes are visualized by the color and size of the dots, respectively.

We also identified the most important genes in the classification of LAEA SIV-positive spots where we found that BSG –also in the top 5 in the regression model–was the most informative gene (**Figure 5C**) closely followed by ATP6. However, these genes were also highly informative in identifying SIV-neighbor spots, possibly indicating their relevance as viral microenvironment markers (**Figure 5B**). Subsequently, the top genes uniquely seen as key features identifying LAEA spots that were most specifically associated with SIV antigen were identified. We selected these genes among the features included in the 75^th^ percentile using SIV classification model importance (top 10 shown in **Figure 5E**). Overall, these genes were associated with metabolic activation, inflammatory microenvironment, innate immunity, migratory capacity and stress responses. Some of the genes with the most notable functions were the following: 1) Two carbonic anhydrases, CA2 –with the highest importance specific to LAEA SIV-positive spots–and CA12, both of which are each highly associated with hypoxia and tumor microenvironments. 2) Multiple genes associated with cytoskeletal dynamics and cell migration, including CDC42, ARHGDIA, IQGAP1, TUBA4A, ACTR3, DBNL, ANXA2. 3) One of the key regulators of the integrated stress response (ISR), ATF4, as well as ER-stress related genes including PRDX6. 4) Genes associated with cancer and tumor metabolism including CA12, SLC16A1 (MCT1), HDGF, CD151, ANXA2, and SEMA3B.

In summary, this analysis identified key markers specifically associated with SIV production during the eclipse phase of an ATI prior to viremia in LAEA (persistent) viral reservoirs. Many of the genes found are highly interconnected and indicate that the persistent reservoirs in LAEA are characterized by activation of stress responses and the establishment of a viral microenvironment that displays features shared with cancer-related processes.

### IF analysis confirms link between SIV presence and ISR key markers

In order to validate the association between SIV foci in gut-tissue reservoirs and the specific activation of ISR in persistent reservoirs, we used IF detection of phosphorylated eukaryotic initiation factor-2α (p-eIF2α), a critical component of the ISR that acts as a switch to reduce global cap-dependent initiation of protein synthesis during cellular stress, in comparison with SIV IF signal. An example of a LAEA tissue stained in this manner is shown in **Figure 6** which reveals two distinct SIV-infected populations (Gag^+^/p-eIF2α^+^ and Gag^+^/p-eIF2α^-^; zoomed-in inserts that show examples of both populations **Figure 6C and E**) in concordance with the two distinct transcriptional patterns attained in the spatial transcriptomics analyses. This observation of a population of p-eIF2α and SIV double positive cells associated with the tissue resident foci of virus replication characterized here, confirms the direct association between SIV infection and ISR activation.

**Fig 6.**
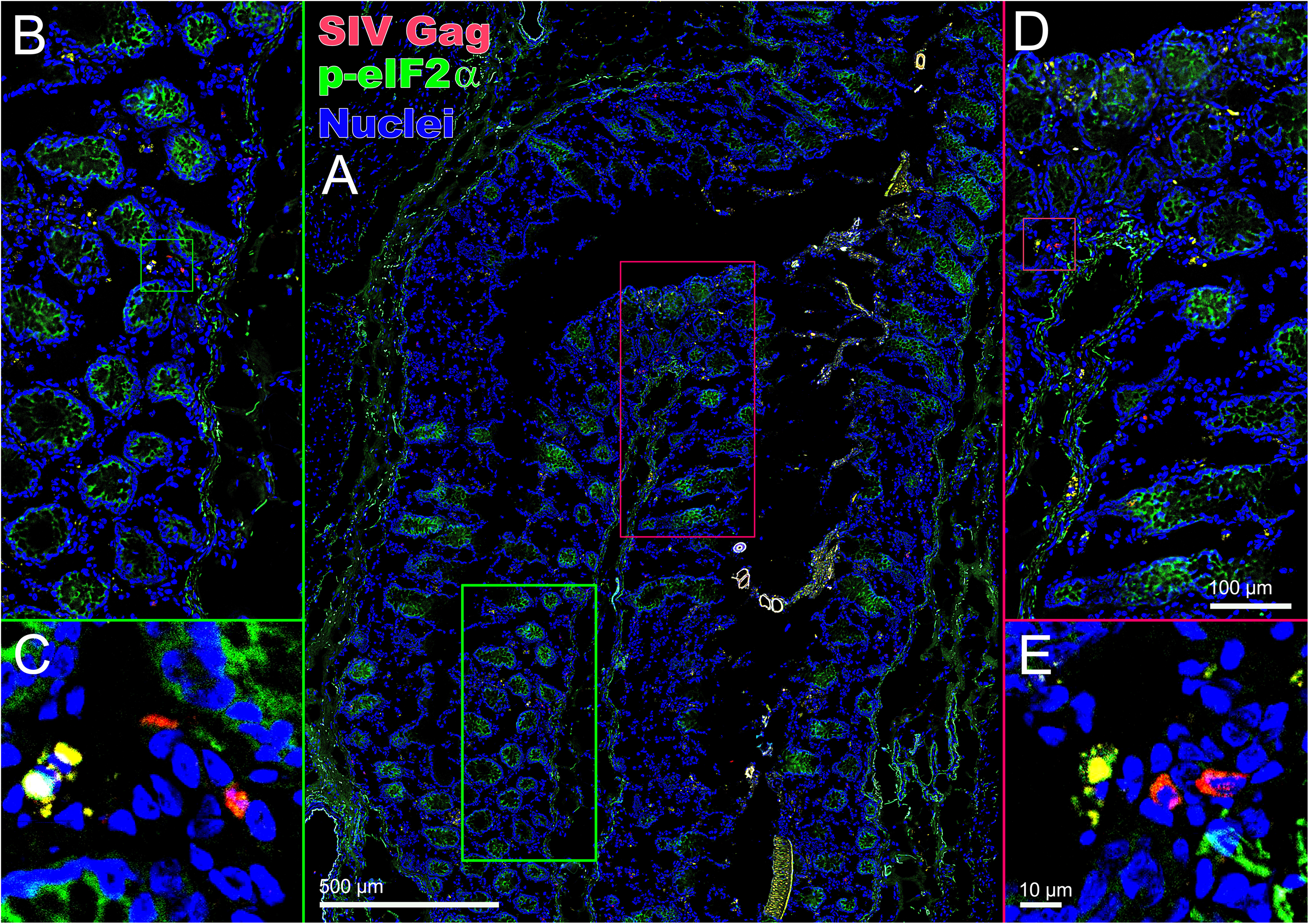
IF validation of SIV association with ISR using ISR marker p-eIF2α. **A)** Representative IF merged image from colon section from LAEA animal (A7005). IF detection of SIV Gag is shown in red, p-eIF2α in green, and cell nuclei in blue. **B and D)** Inserts show two regions respectively of the slide as representative examples of areas where SIV was detected. **C and E)** Detailed imaging within the B and D regions where two populations of SIV Gag+ cells were detected in tissue resident persistent reservoirs in colon tissue; Gag+ p-eIF2α+ in yellow color indicating co-localization and Gag+ p-eIF2α- in red.

## DISCUSSION

This study identifies a specific viral microenvironment (VME) associated with gut tissue resident persistent reservoirs in the SIV-rhesus macaque as a spatially organized tissue niche. Like the analogous TME, the VME functions to sustain viral persistence in spite of suppressive ART. The use of ART efficiently prevents the systemic spread of the virus but does not directly impact the remaining infected cells within the VME. The formation of this VME requires approximately a week to develop after the initial acquisition of infection. By combining immunoPET/CT-guided necropsy with spatial transcriptomics, we directly interrogated tissue resident SIV reservoir sites with foci of viral production during the eclipse phase of rebound. This ability to characterize infected cells during the initial phase of the rebound revealed that reservoir durability is governed by specialized VME features that shape cellular state, immune architecture, and viral production.

Across all animals, viral protein detection was spatially clustered and associated with increased local transcriptional levels, indicating that early rebound in tissues originates from discrete foci rather than stochastic reactivation throughout a tissue. Persistent reservoirs exhibited overall higher SIV foci and proviral burden while displaying lower SIV density values per SIV-positive spots. This is consistent with VMEs that are more deeply entrenched in LAEA reservoir than those in EAEA tissues but are able to produce less virions per infection foci given their reduced translation activity associated with the inhibition of global cap-dependent initiation of translation, a defining aspect of the ISR. Importantly, elevated host transcription was tightly coupled to viral antigen production and not explained by tissue structure alone, positioning the VME as a permissive transcriptional niche that actively supports residual viral expression during ART and initiates early rebound post-ATI.

A defining feature of persistent reservoirs identified here was the activation of cell cycle, metabolic, mitochondrial, and stress-response programs accompanied by repression of global cytoplasmic translation and increased cell senescence. This combination of proliferative signaling, metabolic rewiring, and translational restraint mirrors cellular states observed in tumor and chronic inflammatory microenvironments. Such stress-adapted states may enable long-term survival of infected cells while permitting episodic viral production to circumvent immune and metabolic pressure. In contrast, more transient reservoirs seeded under early ART initiation displayed robust activation of both cytoplasmic and mitochondrial translation, consistent with acutely activated and less constrained cellular programs. These differences indicate that VME maturation over a longer period of untreated infection—rather than viral presence alone—underlies the long-term stability of persistent reservoirs. Additionally, both transient and persistent reservoirs were preferentially localized to luminal gut regions, marked by enrichment of epithelial cells and exclusion of mesenchymal programs. Upregulation of extracellular matrix organization and adhesion pathways in regions outside SIV production enriched areas further supports a model in which viral persistence is favored within structurally remodeled tissue compartments. Persistent reservoirs additionally showed associations with vascular and neuroimmune features, suggesting deeper integration into tissue physiology.

Regarding the cellular components of this VME, our results showed that persistent reservoirs were embedded within immune landscapes resembling immunosuppressed TLS, which we inferred to have a core of either cycling B-cells and Tcm or follicular B-cells and cycling DCs, surrounded by mast cells and ILC, with Treg cells on the periphery. Also, the RBC abundance reflects vascular disruption that co-localize with inflammatory infiltrates and increased SIV antigen density in persistent reservoirs. These environments could foster immune cell exclusion leading to impaired host defense functionality —features compatible with long-term viral persistence. By contrast, transient reservoirs were associated with adaptive effector populations including CD8⁺ T, Th17, Tfh, and activated CD4⁺ T-cells, consistent with immune-active environments capable of rapid antiviral responses. Thus, early ART appears to prevent the consolidation of regulatory VMEs that shield infected cells from immune clearance. Moreover, cell–cell interaction analyses revealed that persistent reservoirs establish a VME orchestrated by Treg-centered communication networks linking SIV-positive, neighboring, and virus-negative regions, with key contributions from ILCs, mast cells, and NK cells. This regulatory wiring likely constrains antiviral effector function while allowing controlled viral reactivation. In contrast, tissue microenvironments associated with transient reservoirs were dominated by Tfh- and Th17-centered interaction networks largely confined to virus-negative and neighboring regions, with minimal immune crosstalk involving SIV-producing sites. These patterns suggest that persistent reservoirs lead to active reprograming of immune cell communication to favor viral persistence.

SIV presence associated to persistent reservoirs was specifically linked to two significantly distinct transcriptional clusters that were also inferred to be comprised by different cellular structures –LAEA-1 and LAEA-6. This is indicative of two different viral populations comprising the detected SIV foci: the reservoir that persisted during ART and the virus that is starting to spread locally during the eclipse phase of the rebound. The former potentially would be mostly associated with LAEA-1, characterized by activation of cell cycle, metabolism, morphogenesis, and host factors that interact with HIV-1, while the latter would be related to LAEA-6, with immune activation signals concordant with newly infecting virus. This observation is also supported by the IF validation where two distinct populations emerge: SIV+p-eIF2α+ and SIV+p-eIF2α-.

The observation of two different clusters associated with SIV-positive spots in both EAEA and LAEA tissues (**Figure 2**) is consistent with the expected pattern of two post ATI tissue resident populations: (1) newly infected cells that expand and disseminate exponentially, and (2) a stable reservoir population that should be rapidly displaced by the expanding footprint of newly infected cells and the associated tissue damage. The integrated analysis of SIV associated clusters (**Figure 2F**) also identified two populations. Notably, the Combined 3 cluster showed an uneven distribution of SIV positive spots, with greater abundance in LAEA, where this cluster also exhibited higher frequencies of SIV positive and SIV neighbor spots. These findings are aligned with emerging immunofluorescence data from the same tissue block in one animal, which revealed two populations of Gag positive cells distinguished by p eIF2α expression (**Figure 6**). The detection of these populations across both spatial transcriptomics and immunofluorescence analyses highlights the strength and rigor of integrating these complementary approaches on adjacent tissue sections.

Finally, unbiased machine learning identified stress-response, metabolic, mitochondrial, cytoskeletal, and hypoxia-associated genes as dominant predictors of viral density and presence within persistent VMEs. Many of these genes—such as carbonic anhydrases, CD147 (BSG), and regulators of cell migration and integrated stress response—are hallmark features of tumor microenvironments. Their convergence across analytical approaches reinforces the concept that persistent VMEs represent a conserved tissue state optimized for survival under stress, immune modulation, and episodic viral production. Moreover, when using IF to validate the co-localization between SIV foci and p-eIF2α, we can clearly observe that this is widely detected in persistent reservoirs, but it is less abundant in transient reservoirs. This orthogonal validation confirms that the signals observed at the transcriptional level are also observed at the protein level. It is also consistent with the rapid, homogeneous rebound associated with persistent reservoirs (LAEA), compared to the heterogenous, delayed rebound characteristic of early ART initiation with ATI after 6 months.

At the end of this study, we identified a set of biomarkers that define the VME specifically associated with persistent HIV-1 reservoirs. To place our findings in the context of existing literature, we compared these VME-defining genes with results from previous studies. Notably, many of the drivers identified here have also been reported elsewhere. For example, we curated genes associated with HIV-1 reservoirs from studies analyzing gut tissue (Wei et al.^35^) and peripheral blood (Clark et al.^36^) in individuals on suppressive therapy. This resulted in a set of 71 unique genes that were significantly upregulated across multiple comparisons: (1) HIV-infected T-cells from gut versus matched HIV-infected T-cells from blood, (2) HIV-infected gut T-cells versus gut T-cells from uninfected donors, and (3) HIV-infected blood CD4⁺ T-cells versus uninfected blood CD4⁺ T-cells. When this curated set was compared with our machine-learning–derived VME-defining gene list (n = 1,282), we observed that 43 of the 71 previously reported genes overlapped with our results, despite the prior studies being restricted to T-cell populations (**Supp. Fig. 14**). Interestingly, the highest overlaps were found between genes differentially upregulated in HIV-infected T-cells compared with uninfected cells both in gut and blood. These similarities underscore the broad relevance of our results and support the applicability of the VME paradigm to HIV-1 persistence. Moreover, they highlight how the contextual and expanded resolution afforded by immunoPET/CT-guided spatial transcriptomics enables a more comprehensive and integrated view of these specialized viral microenvironments, which has not been achievable with prior approaches.

Together, these findings position the VME—not virus transcriptional latency alone—as a central determinant of SIV reservoir durability and rebound off-ART in the non-human primate model. Like an aggressive tumor that has evolved to utilize the TME, a host response which typically controls wayward cells, the virus has evolved to utilize the VME as a sanctuary. Persistent reservoirs are stabilized by VMEs that integrate stress adaptation, immune regulation, and tissue remodeling. Consequently, latency reversal strategies may fail unless paired with interventions that dismantle the VME itself. Targeting translational repression, senescence, metabolic rewiring, or Treg-mediated immune control may be required to destabilize persistent VMEs and enable immune-mediated clearance.

## METHODS

### Ethical statement

All animal studies were conducted in accordance with protocols approved by Northwestern University and New Iberia Research Center (NIRC), University of Louisiana at Lafayette Local Institutional Animal Care and Use Committees (IACUCs), protocol number 2017-8791-002. This study was carried out in accordance with the Guide for the Care and Use of Laboratory Animals of the NIH and with the recommendations of the Weatherall report ‘‘The use of nonhuman primates in research.’’ All procedures were performed under anesthesia using ketamine hydrochloride (10mg/kg) and/or Telazol (4mg/kg), and all efforts were made to minimize stress, improve housing conditions, and provide enrichment opportunities. Animals were euthanized by sedation with ketamine hydrochloride injection followed by intravenous barbiturate overdose in accordance with the recommendations of the panel on euthanasia of the American Veterinary Medical Association.

### Animals

A total of 7 adult male and female rhesus macaques (*Macaca mulatta*) [RM] were sourced from the NIRC colonies (**Supp. Table 3**). All animals were determined to be specific pathogen free, as defined by being free of cercopithecine herpesvirus 1, D-type simian retrovirus, simian T-lymphotropic virus type 1 and *Mycobacterium tuberculosis*. Additionally, they were typed for protective MHC alleles MHC MamuA*001, B*008 and B*017, as well as for TRIM5α and FcRy to avoid restrictive genotypes. For the entire duration of the study, the animals were housed in the BSL-2+ containment facility at the NIRC and monitored daily for clinical appearance and general health status. Two groups of animals were compared (**Supp. Figure 1A**): Early ART / Early ATI [EAEA] (n=4), and Late ART / Early ATI [LAEA] (n=3). To model the reservoir with a short lifespan (“transient” reservoir) established quickly after SIV acquisition, RMs from the EAEA group that were challenged intravaginally/intrarectally with a 2000 TCID_50_ dose of either SIVmac239, or SIV-ffLuc^37^ virus were initiated on ART at 4 days post-infection. Animals in the EAEA group were maintained under viremia-suppressing ART for 6 months followed by ATI and euthanized 4-5 days after ATI started (during the eclipse phase before viremia). On the other hand, to model a “persistent” reservoir, the LAEA group RMs were started on viremia-suppressing ART at week 10 after intra-venous challenge with the barcoded virus SIVmac239M2 with a 300 TCID_50_ dose and maintained on ART for 54 weeks before an ATI. The 4 EAEA RMs were euthanized 4-5 days post-ATI when viral load (VL) was below detection. The LAEA RMs were euthanized 5-6 days post-ATI when viral load (VL) was below detection in 2 of 3 animals. One LAEA animal had plasma VL=1.2×10^−3^ copies/mL at time of euthanasia. For all animals, plasma viral RNA was regularly tracked throughout the entire course of ART-treated infection and ATI. While RM in the EAEA group never showed consistently detectable plasma viral RNA, just blips. Those in the LAEA group showed a typical slowly declining viremia after ART initiation **(Supp. Figure 1B**). For this latter group, 2 viral blips were also induced by therapy cessation for 3 days before ART re-initiation at weeks 22 and 35 post-ART to ensure robust reservoir seeding. All animals were treated with combined ART corresponding to a human first line triple therapy comprising tenofovir (20 mg/kg/day subQ), emtricitabine (40 mg/kg/day subQ) and an integrase inhibitor, either Dolutegravir, 2.5 mg/kd/day subQ or Raltegravir 100 mg/day orally.

### ImmunoPET/CT

Foci of productively infected cells were identified in vivo with immunoPET/CT using a ^64^Copper-labelled F(ab)2 of the primatized 7D3 antibody to SIV gp120^38,39^. On the day of euthanasia and 18-24 hours after intravenous ^64^Cu-labelled 7D3 probe injection, a whole-body PET-CT scan was performed (**Supp. Figure 1C**) to identify organs and areas with detectable SIV production foci. The identified gut regions were carefully isolated at necropsy as “hot” tissue areas and subjected to a second PET-CT scan to confirm their signal. Tissue regions containing positive probe signal were then cut into smaller blocks, placed in cryomolds and rescanned with immunoPET/CT to identify individual “hot” tissue blocks (**Supp. Figure 1C**). Gut-tissue blocks selected for spatial transcriptomics were serially sectioned to attain tissue sections axially separated by 20µm for spatial transcriptomics analysis, proviral Quantitative Real-Time PCR (qPCR) or digital PCR (dPCR) and for immunofluorescent (IF) staining with antibodies against SIV Gag and immune cell markers.

### Quantification of SIV proviral DNA and viral RNA

Tissue associated proviral DNA and viral RNA were extracted using AllPrep DNA/RNA Kit (Qiagen) per the manufacturer’s protocol. From viral RNA, cDNA was synthesized utilizing random hexamers and the Superscript IV Synthesis Kit (ThermoFisher). Copy number quantification was then performed via qPCR targeting SIV gag using PrimeTime Gene Expression Master Mix (Integrated DNA Technologies). In brief, sample DNA or cDNA was combined with forward and reverse primers (500nM each) and a probe (250nm) targeting SIV gag (SIV gag F: 5′-GTCTGCGTCATYTGGTGCATTC-3′; SIV gag R: 5’-CACTAGRTGTCTCTGCACTATYTGTTTTG-3’; SIV gag Probe: 5′-FAM-CTTCYTCAGTRTGTTTCACTTTCTCTTCTGCG-MGB-NFQ-3′), 60x primer-limited RPL32 Gene Expression Assay Rh02811772_s1 (ThermoFisher), nuclease-free water, and 2x PrimeTime for a total reaction volume of 20µL. Reactions were performed in duplicate following the manufacturer’s suggested cycling conditions. Standard curves for SIV gag and Rh RPL32 were generated using a 5-fold serial dilution of purified 3D8 cells (RRID:CVCL_U276) DNA and Rhesus Monkey control genomic DNA (Fisher Scientific) to allow for validation of amplification efficiencies and determine SIV gag and host copy numbers. Samples with low copy numbers or no amplification in qPCR were confirmed via the QuantStudio Absolute Q Digital PCR system (Applied Biosystems). Reaction mixtures had a final volume of 10µL, consisting of sample DNA or cDNA, SIV gag primers/probe, Rh RPL32 assay, nuclease-free water, and 5x Absolute Q DNA Digital PCR Master Mix (ThermoFisher). All runs included a positive control and a non-template control. Reactions were run as per the manufacturer’s cycling conditions. After fluorescence thresholding, the absolute SIV gag and host copy numbers are determined in copies per µL.

### Tissue Immunostaining and Microscopy

IF staining of frozen gut tissues embedded in OCT was conducted according to previously published methods^40^. Briefly, 10 µm thick tissue sections were fixed with 4% formaldehyde in PIPES [piperazine-N, N’-bis(2-ethanesulfonic acid)] buffer (Sigma) and blocked with donkey serum to prevent non-specific antibody binding. Infected cells were identified by staining with mouse anti-SIV Gag (Ag3.0, hybridoma supernatant^41^). Other antibodies used for the study were: mouse phosphorylated eIF2α (1A4A11, ThermoFisher), CD3 (SP7, Abcam), CD117/c-Kit (104D2, Biolegend), and Hoescht nuclear stain. Images were acquired on DeltaVision inverted microscope as four color image stacks containing 3 sections in the Z-plane, projected, and deconvolved using softWoRx software (Applied Precision). Images were analyzed with softWoRx software (Applied Precision) and QuPath v0.3.2 (4).

### Spatial transcriptomics of necropsied gut tissues

Gut-tissue sections identified as “hot” sections by PET/CT-guided necropsy were selected for spatial transcriptomics analysis after RNA quality assessment using TapeStation RNA ScreenTape. Four tissue sections were placed on each of the 10x Visium Spatial Gene Expression Slides (10X Genomics), fixed with chilled methanol, H&E stained and imaged under a brightfield microscope. Afterwards, reverse transcription and barcoding was performed *in situ* and libraries were prepared according to the Visium Spatial Gene Expression Slide & Reagent Kit (PN-1000184) according to the manufacturer’s protocol (CG000239). Generated libraries were subsequently sequenced in NUSeq Core on the Illumina Novaseq 6000 platform. Sequences and histology images were subsequently processed with the Space Ranger software v.1.3.1, using the macaque reference genome Mmul10. Visualization, integration, and cell clustering and differential expression analyses were performed using the R package Seurat v.5.1.047^42^.

### Spatial transcriptomics Analysis

Raw counts were quality filtered and subjected to Principal Component Analysis (PCA)-based dimensionality reduction and Louvain clustering at 0.3 resolution using the 15 most informative principal components; clusters were visualized using UMAP and t-SNE. Spatially variable genes were identified using Moran’s I per slide. For combined analyses, slides were integrated per group using the harmony pipeline^43^ in Seurat and identified differentially expressed genes (DEG) and spatial enrichment for each transcriptional cluster. DEG analyses were performed after raw count normalization with SCTransform using statistical modeling using the Model-based Analysis of Single-cell Transcriptomics (MAST) framework^44^ controlling for the slide of origin of each spot.

After initial transcriptional clustering and differential expression analysis independently of SIV antigen detection, we identified clusters and their corresponding markers differentially expressed in SIV Gag-positive versus -negative spots using the same DEG approach in pairwise group comparisons. To define such SIV-positive spots, we associated 10X Visium Spatial Gene Expression Slide spots coordinates with the location coordinates of SIV Gag antigen detected by IF in axially adjacent tissue sections separated by 20µm from the one mounted in the 10X Slide (**Supp. Figure 1D**) using SPATA2 v.2.0.329^45^ and in-house R scripts. The criterion to classify a spot as containing SIV-infected cells was if SIV Gag detected by IF was located within 50µm distance from the centroid of the Visium spot in any direction, covering this way the ideal 3D space enclosed within the boundaries of the spots (diameter=55µm) plus the average space between the center of the spots (100µm). Following this approach we created two spot categories: “SIV-Positive” and “SIV-Negative” (**Supp. Figure 1D**). For more detailed analyses, we also added an additional spot tagging strategy where we assigned the neighboring spots within 100 µm to the centroids of the ones previously tagged as SIV-positive, as “SIV-Neighbors”. We then performed the same analysis but now comparing the three groups of spots “SIV-Positive”, “SIV-Neighbor”, and “SIV-Negative”. After DEG analysis, we performed gene set enrichment analysis (GSEA)^46^ using the canonical pathways subcollection and the Hallmark gene sets included in the macaque Molecular Signatures Database (MSigDB) and the macaque Gene Ontology (GO) database. For such analyses, we used all the transcripts that appeared in at least 1% of the analyzed spots ranked by the log-fold change between SIV-Positive and SIV-Negative spots or additional corresponding group comparisons (e.g., direct pairwise cluster comparisons or SIV-positive vs SIV-neighbor comparisons).

### Validation of Tissue Preparation and spatial transcriptomics Analytical Methods

After optimizing immunoPET/CT guided resection, embedding of “hot” tissues, sectioning, permeabilization, and slide coverage maximization, spatial transcriptomics sensitivity and specificity was validated using a single slide from a transverse colon necropsied at 4 days post-ATI start from an EAEA animal. This began with IF detection of SIV Gag, CD3 and CD117 positive cells in an axially adjacent tissue section separated by 20µm. The IF analysis allowed validating the SIV Env immunoPET/CT-guided necropsy and confirmed preservation of structures between adjacent tissue slices to enable their complementary analysis.

In the transverse colon section used for validation, IF and H&E microscopy validated that cell antigen expression patterns were consistent with the spatial transcriptomics findings, allowing localization in sections of SIV antigen producing cells as well as other immune cells (**Supp. Figure 2A**). Comparison of gene expression clusters with microscopy identified an immune aggregate visible on H&E imaging (indicated with purple circles and lines in **Supp. Figure 2A**) that had a high frequency of CD3+ cells and very rare detection of SIV Gag antigen producing cells by IF. This immune aggregate completely coincided with a single transcriptional cluster (cluster 6). (**Supp. Figure 2B and C**). Further, the genes that were significantly differentially expressed in this cluster compared to the other clusters were highly enriched with genes associated with GO terms related to immune processes (**Supp. Figure 2D**), including “positive regulation of immune system process”, “lymphocyte activation”, “B cell receptor signaling pathway”, “adaptive immune response”, or “antigen processing and presentation of peptide or polysaccharide antigen via MHC class II.” Similarly, an area displaying high density of SIV Gag and CD117 staining (indicated by red circles and lines in **Figure 2A**) overlapped with cluster 3. This cluster was characterized by differential enrichment for GO terms associated with viral production such as “viral process”, “viral release from host cell”, or “exit from host cell” (**Supp. Figure 2E**). Complementary enrichment analysis using REACTOME was done only for this validation and also showed HIV infection as one of the top enriched pathways in this cluster.

### Ortholog conversion of rhesus macaque genes to human genes

For multiple analyses human datasets were used, thus ortholog conversion between RM and human was necessary. To perform this ortholog conversion, all gene names present in the spatial transcriptomics anndata object within the anndata.var layer were checked against a dictionary of *Macaca mulatta*/*Homo sapiens* gene orthologs generated using the biomarRt package in R. The numpy function numpy.intersect1d in Python was used to determine which genes in the spatial transcriptomics data have corresponding rhesus macaque orthologs. Upon determining which genes had orthologs in the dictionary, the numpy function numpy.where was used to select indices of the gene names at which there was a RM gene ortholog to be converted to a human gene. At indices where an orthologous human gene name was available, the rhesus macaque gene name was replaced, in place at the corresponding index, to the human ortholog. Genes that mapped to multiple orthologs were not converted to prevent duplicated gene names.

### Senescence Scoring

Senescence scoring was completed using the SenePy package in Python^32^. The hub settings species= “human” and hubs.metadata.tissue= “intestine” were used as backgrounds for senescence scoring. The hubs.get_genes function was used to select the categories “intestine” and “epithelial cell” for scoring. Raw spatial transcriptomics data for the EAEA and LAEA conditions were concatenated into a single anndata object and preprocessed (e.g., filtration, normalization, etc.) using standard scanpy functions (scanpy.preprocessing.normalize, scanpy.preprocessing.log1p, scanpy.preprocessing.pca, scanpy.preprocessing.neighbors, scanpy.tools.umap, scanpy.tools.leiden). Gene names in spatial transcriptomics data were converted from rhesus macaque to human orthologs and the translator function in SenePy was used to maximize spatial transcriptomics gene overlap with the SenePy built-in human gene expression reference used. Senescence scores were then acquired using the SenePy command senepy.score_hub and scores were added to the anndata.obs metadata for visualization. Senescence scoring analysis was completed in Python 3.10.18, using SenePy version 1.0.1.

### Spatial enrichment

Spatial enrichment analysis was completed using the decoupleR package in Python^47^. Raw data was read into Python and concatenated within each experimental condition (EAEA vs. LAEA) and normalized using standard scanpy functions (scanpy.preprocessing.normalize, scanpy.preprocessing.log1p). Labels for SIV group (SIV-negative, SIV-neighbor, and SIV-positive) were added to the metadata by spatial barcode. Gene sets from the GO biological processes (BP) category in the MSigDB pathway lists were loaded into Python and scored for spatial enrichment using the decoupler.pp.get_obsm function to obtain a composite score for each pathway at each spatial transcriptomics spot. Several t-tests were then completed with FDR correction stratifying by SIV-negative, SIV-neighbor, and SIV-positive categories to determine which gene pathways were spatially enriched among each of these categories in comparison to all other spots on the tissue. The function decoupler.tools.rankby_group was used to generate a data frame of spatial enrichments by SIV category. Spatial enrichment analysis was completed in Python 3.10.19, using decoupleR version 2.1.2.

### Cell type frequency inference per spot

Cell type proportions within spatial transcriptomics spots were identified using the Deconvolution of Spatial Transcriptomics profiles using Variational Inference (DestVI) package in Python^48^. Two single cell RNA sequencing (scRNA-seq) references were used to deconvolute cell types: one scRNA-seq dataset from macaques (*M. fascicularis*) as an initial reference^49^ and a human reference from the Space-Time Gut Cell Atlas within the Human Cell Atlas^50^. A subset of the top 2500 highly variable genes was selected from the scRNA-seq data, followed by standard preprocessing of the scRNA-seq data using scanpy functions^51^ (scanpy.preprocessing.filter_genes, scanpy.preprocessing.highly_variable_genes, scanpy.preprocessing.normalize, and scanpy.preprocessing.log1p). Raw spatial transcriptomics data for each animal group were read into Python as anndata objects, concatenated among slides within the respective condition and preprocessed, also using standard scanpy functions. For cell type identification using the human scRNA-seq reference background, preprocessing included converting gene names in spatial transcriptomics data from RM to human orthologs, as described in the *Ortholog conversion of rhesus macaque genes to human genes* section. A subset of the preprocessed scRNA-seq and preprocessed spatial transcriptomics data was created based on intersecting genes present in both datasets (using the numpy.intersect1d function in Python to find overlapping genes between the scRNA-seq and spatial transcriptomics data). The scRNA-seq data was used to build and train a machine learning model as a reference background for cell type deconvolution of the spatial transcriptomics data. Upon identification of cell type frequencies in spots on the tissue slides, a correlation analysis was completed using the “corr” function in R to calculate the Spearman correlation between the rhesus macaque and human cell type identification frequencies to validate the utility of the spatial transcriptomics cell type identification based on the human scRNA-seq reference for further analyses. A strong positive correlation was identified among most cell types between the inferred cell type frequencies from the macaque scRNA-seq background and human scRNA-seq background (**Supp. Figure 15**). Consequently, the inferred cell type proportions acquired from the human scRNA-seq background were used for further analyses. Given the rich immune microenvironment of the gut, a more granular analysis of immune cell types present in the tissues was completed by running another DestVI cell type deconvolution analysis only on barcodes for which at least 1% of cells had the cell type label “B cells”, “Myeloid”, “Neuronal”, “Plasma cells”, “Red blood cells”, and “T cells” using a reference background from the Colon Cell Atlas scRNA-seq database^52^ that contains a more diverse array of immune cell subtypes present in the gut microenvironment. For all cell types present in the analysis refer to **Supp. Table 4**) Analysis was completed with Python 3.12.10 and scvi-tools version 1.3.0.

### Cell-cell interactions

Cell-cell interactions (CCIs) were determined through ligand-receptor-based analysis using the stLearn package in Python^53^. Raw spatial transcriptomics data files were read into Python and concatenated within each animal group (EAEA vs. LAEA) into an anndata object. Anndata objects for each condition were preprocessed using standard stLearn functions (stlearn.preprocessing.filter_genes, stlearn. preprocessing.normalize_total). Metadata from spots labeled as Neg, SIV neighbor, and SIV were merged with cell type labels inferred from the DestVI analysis using spatial barcodes to create new combined labels (i.e., Treg_SIVneighbor). Cell type frequencies for all cell types at each barcode were added to the anndata.uns layer of the anndata object, and the maximum cell type frequency label for each barcode was added to the anndata.obs data layer. Of note, in the matrix of cell type proportions, for each of the SIV categories (SIV-negative, SIV-neighbor, and SIV-positive), cell type frequencies were labeled as 0% for the two conditions that were not applicable at the respective spot (i.e., for SIV spots, cell type frequencies for the SIV-negative and SIV-neighbor columns were labeled as 0%). The connectomeDB2020_lit database was then used to acquire reference data on ligand-receptor interactions, using the ‘macaque’ species setting. Gene names beginning with ‘Metazoa’, ‘Y’, ‘5S’, ‘RN’, ‘5_8’ were filtered out of the spatial transcriptomics dataset for ligand-receptor interaction identification. Ligand-receptor analysis was then run to determine statistically significant cell-cell interactions using the stLearn commands stlearn.tools.cci_run (settings: min_spots=20, n_pairs=10,000) and stlearn.tools.cci.run_cci (settings min_spots=3, spot_mixtures=True, cell_prop_cutoff=0.05, sig_spots=True, n_perms=1000). Analysis was completed using Python version 3.10.17 and stLearn version 0.4.11.

### Principal component analysis

Principal component analysis (PCA) was performed on each of the inferred cell type datasets (FactoMineR v2.13 package) with scaling, retaining 5 components, and variable contributions were examined per axis. Hierarchical clustering on principal components (HCPC) was then applied to the PCA output to determine the optimal number of clusters automatically, and resulting cluster assignments were merged back into the spot metadata for downstream visualization and comparison with Seurat clusters and SIV regions.

### Machine Learning Classification

Multiclass classification was performed using the mlr3 framework^54^ v.1.5.0 in R to predict group membership from viral raw count data. The top 1000 features were selected using information gain filtering. Five algorithms were evaluated via 3-fold cross-validation: XGBoost, Random Forest (Ranger), LightGBM, Elastic Net (GLMnet), and Support Vector Machine (SVM). The best-performing model, selected based on multiclass log loss, underwent Bayesian hyperparameter optimization using Model-Based Optimization (MBO) with 25 evaluations and 3-fold cross-validation. Model performance was assessed using classification accuracy, log loss, and Brier score, with confusion matrices generated to visualize classification patterns. Feature importance was extracted from the optimized model, and class-specific importance was calculated for each region (SIV-positive, SIV-neighbor, and SIV-negative) and group (EAEA and LAEA) using binary classification (80/20 train-test split). Expression patterns for top features were visualized using ggbeeswarm v.0.7.3 R package plots. All analyses used a random seed of 123 with 15-worker parallel processing for reproducibility and computational efficiency.

### Machine Learning Regression Analysis

Regression analysis was also performed using the mlr3 framework in R to predict SIV density from viral raw count data. The top 1000 features were selected using variance-based filtering. Five algorithms were evaluated via 3-fold cross-validation: Random Forest (Ranger) with permutation importance, XGBoost, Linear Regression, Support Vector Machine (SVM), and LightGBM. All models except linear regression underwent Bayesian hyperparameter optimization using MBO with 25 evaluations and holdout resampling, optimizing for R-squared. The best-performing model, selected based on lowest RMSE, was further validated using 5-fold cross-validation and subsequently trained on 80% of the data with final evaluation on the remaining 20% test set. Performance was assessed using RMSE, MAE, R-squared, and MSE. Feature importance was extracted using the model’s native method (permutation-based for Random Forest), and the top 50 features were visualized with numeric scores. Model predictions were evaluated using predicted versus actual plots with a reference line. All analyses used a random seed of 123 with 15-worker parallel processing for reproducibility and computational efficiency.

### Statistical modeling

All statistical modeling analyses were performed in R v4.4.1. Simple group comparisons were performed using Wilcoxon Rank Sum tests. Overall transcriptional counts or SIV density values were summarized across SIV regions and experimental group, then modeled using negative binomial generalized linear mixed models (glmer.nb function from lme4 v1.1-38 package) with slide identity as a random effect. The effect of SIV-positive vs. SIV-negative and its interaction with group on median transcript counts was assessed in one model, while a second model evaluated differences between the three SIV regions (SIV-positive, SIV-neighbor, and SIV-negative). Pairwise comparisons were derived using estimated marginal means (emmeans package) with FDR correction. Changes in SIV density per group for all SIV-positive spots were tested using a generalized linear model with a Gamma distribution and log link controlling for time since ATI of the sample. Spatial clustering of SIV-positive spots was assessed using a quadrat test with Monte Carlo simulation (n = 5,000 iterations) implemented via the spatstat v3.5-1 package in R, where cell centroids were modeled as a planar point process and tested against complete spatial randomness.

Finally, for cell type analysis cell types present in at least 10% of spots were retained. Two complementary approaches were used to assess the relationship between immune cell composition and SIV presence. For the continuous approach, the association between cell type frequency and SIV density was modeled for each cell type independently using linear mixed models (lme4), with log-transformed SIV density as the outcome, cell type frequency and log-transformed total transcript count as fixed effects, and sample identity as a random effect. P-values were corrected for multiple comparisons using the Benjamini-Hochberg method, and cell types with FDR < 0.05 were considered significant. For the categorical approach, differences in cell type frequency between spots from the three different SIV regions were modeled jointly across all cell types using a Tweedie generalized linear mixed model (glmmTMB v1.1.14) with an interaction term between SIV category and cell type, and sample identity as a random effect. Pairwise contrasts were extracted using estimated marginal means with FDR correction.

## Acknowledgements

The authors thank Dr. Elena Martinelli for initial discussions of the spatial transcriptomics approach and coordinating the transfer of plasma from NIRC to Leidos for pVL determination. Thank Dr. Jeff Lifson and Leidos for pVL determination and Dr. Brandon Keele for providing the SIVmac239M2 virus. This research was additionally supported through the computational resources and staff contributions provided by the Quest high-performance computing facility and Sequencing Core facility (NUSeq) at Northwestern University, which is jointly supported by the Office of the Provost, the Office for Research, and Northwestern University Information Technology.

## Data Availability

All scripts and bioinformatic pipelines are available at the Lorenzo-Redondo lab github (https://github.com/rlr-lab/). All processed files and imaging files are stored on the Northwestern FSMResFiles centralized data storage platform for long-term storage. Sequencing files are deposited to NCBI as Bioproject.

## Author contributions

T.J.H, F.V., and R.L-R conceived and designed the study; M.S.A., Y.T., E.Z., C.T.T., R.V.M., F.E., J.M.H., S.H.B., M.A.S, E.J.A., A.M.C., D.F., N.O., M.D.M., M.A., and F.V. performed the experiments; E.U.C., M.S.A., S.C.P., T.J.H and R.L-R. analyzed the data; T.J.H, E.U.C., A.M., I.C., R.T.D., J.F.H., and R.L-R interpreted the results; T.J.H, E.U.C., R.T.D., J.F.H., and R.L-R wrote the paper. All the authors read, reviewed, and approved the final manuscript.

## References

1. Gantner, P., Buranapraditkun, S., Pagliuzza, A., Dufour, C., Pardons, M., Mitchell, J.L., Kroon, E., Sacdalan, C., Tulmethakaan, N., Pinyakorn, S., et al. (2023). HIV rapidly targets a diverse pool of CD4(+) T cells to establish productive and latent infections. Immunity 56, 653–668 e655. 10.1016/j.immuni.2023.01.030.

2. Pieren, D.K.J., Benitez-Martinez, A., and Genesca, M. (2024). Targeting HIV persistence in the tissue. Curr Opin HIV AIDS 19, 69–78. 10.1097/COH.0000000000000836.

3. Chaillon, A., Gianella, S., Dellicour, S., Rawlings, S.A., Schlub, T.E., De Oliveira, M.F., Ignacio, C., Porrachia, M., Vrancken, B., and Smith, D.M. (2020). HIV persists throughout deep tissues with repopulation from multiple anatomical sources. J Clin Invest 130, 1699–1712. 10.1172/JCI134815.

4. Estes, J.D., Kityo, C., Ssali, F., Swainson, L., Makamdop, K.N., Del Prete, G.Q., Deeks, S.G., Luciw, P.A., Chipman, J.G., Beilman, G.J., et al. (2017). Defining total-body AIDS-virus burden with implications for curative strategies. Nat Med 23, 1271–1276. 10.1038/nm.4411.

5. Busman-Sahay, K., Starke, C.E., Nekorchuk, M.D., and Estes, J.D. (2021). Eliminating HIV reservoirs for a cure: the issue is in the tissue. Curr Opin HIV AIDS 16, 200–208. 10.1097/COH.0000000000000688.

6. Cole, B., Lambrechts, L., Boyer, Z., Noppe, Y., De Scheerder, M.A., Eden, J.S., Vrancken, B., Schlub, T.E., McLaughlin, S., Frenkel, L.M., et al. (2022). Extensive characterization of HIV-1 reservoirs reveals links to plasma viremia before and during analytical treatment interruption. Cell Rep 39, 110739. 10.1016/j.celrep.2022.110739.

7. Keele, B.F., Okoye, A.A., Fennessey, C.M., Varco-Merth, B., Immonen, T.T., Kose, E., Conchas, A., Pinkevych, M., Lipkey, L., Newman, L., et al. (2024). Early antiretroviral therapy in SIV-infected rhesus macaques reveals a multiphasic, saturable dynamic accumulation of the rebound competent viral reservoir. PLoS Pathog 20, e1012135. 10.1371/journal.ppat.1012135.

8. Terrade, G., Huot, N., Petitdemange, C., Lazzerini, M., Orta Resendiz, A., Jacquelin, B., and Muller-Trutwin, M. (2021). Interests of the Non-Human Primate Models for HIV Cure Research. Vaccines (Basel) 9. 10.3390/vaccines9090958.

9. Trifone, C., Richard, C., Pagliuzza, A., Dufour, C., Lemieux, A., Clark, N.M., Janaka, S.K., Fennessey, C.M., Keele, B.F., Fromentin, R., et al. (2025). Contribution of intact viral genomes persisting in blood and tissues during ART to plasma viral rebound in SHIV-infected rhesus macaques. iScience 28, 111998. 10.1016/j.isci.2025.111998.

10. Okoye, A.A., Duell, D.D., Fukazawa, Y., Varco-Merth, B., Marenco, A., Behrens, H., Chaunzwa, M., Selseth, A.N., Gilbride, R.M., Shao, J., et al. (2021). CD8+ T cells fail to limit SIV reactivation following ART withdrawal until after viral amplification. J Clin Invest 131. 10.1172/JCI141677.

11. Liu, P.T., Keele, B.F., Abbink, P., Mercado, N.B., Liu, J., Bondzie, E.A., Chandrashekar, A., Borducchi, E.N., Hesselgesser, J., Mish, M., et al. (2020). Origin of rebound virus in chronically SIV-infected Rhesus monkeys following treatment discontinuation. Nat Commun 11, 5412. 10.1038/s41467-020-19254-2.

12. Keele, B.F., Okoye, A.A., Immonen, T.T., Varco-Merth, B., Duell, D., Nkoy, C., Goodwin, W., Hoffmeister, S., Hughes, C.M., Kose, E., et al. (2026). Initial sites of SIV rebound after antiretroviral treatment cessation in rhesus macaques. Nat Microbiol 11, 648–663. 10.1038/s41564-025-02258-3.

13. Gunst, J.D., Gohil, J., Li, J.Z., Bosch, R.J., White Catherine Seamon, A., Chun, T.W., Mothe, B., Gittens, K., Praiss, L., De Scheerder, M.A., et al. (2025). Time to HIV viral rebound and frequency of post-treatment control after analytical interruption of antiretroviral therapy: an individual data-based meta-analysis of 24 prospective studies. Nat Commun 16, 906. 10.1038/s41467-025-56116-1.

14. Weerasuria, M., McMahon, J.H., Lewin, S.R., and Lau, J.S.Y. (2025). The role of analytical treatment interruptions in shaping HIV-specific immunity and HIV cure. Curr Opin HIV AIDS 20, 543–551. 10.1097/COH.0000000000000973.

15. Gondim, M.V.P., Sherrill-Mix, S., Bibollet-Ruche, F., Russell, R.M., Trimboli, S., Smith, A.G., Li, Y., Liu, W., Avitto, A.N., DeVoto, J.C., et al. (2021). Heightened resistance to host type 1 interferons characterizes HIV-1 at transmission and after antiretroviral therapy interruption. Sci Transl Med 13. 10.1126/scitranslmed.abd8179.

16. Bertagnolli, L.N., Varriale, J., Sweet, S., Brockhurst, J., Simonetti, F.R., White, J., Beg, S., Lynn, K., Mounzer, K., Frank, I., et al. (2020). Autologous IgG antibodies block outgrowth of a substantial but variable fraction of viruses in the latent reservoir for HIV-1. Proc Natl Acad Sci U S A 117, 32066–32077. 10.1073/pnas.2020617117.

17. Okoye, A.A., Hansen, S.G., Vaidya, M., Fukazawa, Y., Park, H., Duell, D.M., Lum, R., Hughes, C.M., Ventura, A.B., Ainslie, E., et al. (2018). Early antiretroviral therapy limits SIV reservoir establishment to delay or prevent post-treatment viral rebound. Nat Med 24, 1430–1440. 10.1038/s41591-018-0130-7.

18. Wang, X., Vincent, E., Siddiqui, S., Turnbull, K., Lu, H., Blair, R., Wu, X., Watkins, M., Ziani, W., Shao, J., et al. (2022). Early treatment regimens achieve sustained virologic remission in infant macaques infected with SIV at birth. Nat Commun 13, 4823. 10.1038/s41467-022-32554-z.

19. Passaes, C., Desjardins, D., Chapel, A., Monceaux, V., Lemaitre, J., Melard, A., Perdomo-Celis, F., Planchais, C., Gourves, M., Dimant, N., et al. (2024). Early antiretroviral therapy favors post-treatment SIV control associated with the expansion of enhanced memory CD8(+) T-cells. Nat Commun 15, 178. 10.1038/s41467-023-44389-3.

20. Gutierrez, H., and Eugenin, E.A. (2024). The challenges to detect, quantify, and characterize viral reservoirs in the current antiretroviral era. NeuroImmune Pharm Ther 3, 211–219. 10.1515/nipt-2024-0017.

21. Taylor, R.A., McRaven, M.D., Carias, A.M., Anderson, M.R., Matias, E., Arainga, M., Allen, E.J., Rogers, K.A., Gupta, S., Kulkarni, V., et al. (2021). Localization of infection in neonatal rhesus macaques after oral viral challenge. PLoS Pathog 17, e1009855. 10.1371/journal.ppat.1009855.

22. Taylor, R.A., Xiao, S., Carias, A.M., McRaven, M.D., Thakkar, D.N., Arainga, M., Lorenzo-Redondo, R., Allen, E.J., Rogers, K.A., Kumarapperuma, S.C., et al. (2024). PET/CT Targeted Tissue Sampling Reveals Intravenously Administered HGN194 IgG1 Affects HIV Distribution after Rectal Exposure. AIDS Res Hum Retroviruses 40, 637–648. 10.1089/AID.2024.0019.

23. Samer, S., Thomas, Y., Arainga, M., Carter, C., Shirreff, L.M., Arif, M.S., Avita, J.M., Frank, I., McRaven, M.D., Thuruthiyil, C.T., et al. (2022). Blockade of TGF-beta signaling reactivates HIV-1/SIV reservoirs and immune responses in vivo. JCI Insight 7. 10.1172/jci.insight.162290.

24. Xu, R., Liu, H., Zhu, T., Tang, H., Wu, M., Yan, X., Li, M., Yuan, S., Yin, T., Chen, J., et al. (2026). An immunosuppressive tertiary lymphoid structure is associated with adverse prognosis in gastric-type endocervical adenocarcinoma. J Natl Cancer Inst 118, 276–288. 10.1093/jnci/djaf310.

25. Zhao, L., Jin, S., Wang, S., Zhang, Z., Wang, X., Chen, Z., Wang, X., Huang, S., Zhang, D., and Wu, H. (2024). Tertiary lymphoid structures in diseases: immune mechanisms and therapeutic advances. Signal Transduct Target Ther 9, 225. 10.1038/s41392-024-01947-5.

26. Pakos-Zebrucka, K., Koryga, I., Mnich, K., Ljujic, M., Samali, A., and Gorman, A.M. (2016). The integrated stress response. EMBO Rep 17, 1374–1395. 10.15252/embr.201642195.

27. Ryoo, H.D., and Vasudevan, D. (2017). Two distinct nodes of translational inhibition in the Integrated Stress Response. BMB Rep 50, 539–545. 10.5483/bmbrep.2017.50.11.157.

28. Wek, R.C., Anthony, T.G., and Staschke, K.A. (2023). Surviving and Adapting to Stress: Translational Control and the Integrated Stress Response. Antioxid Redox Signal 39, 351–373. 10.1089/ars.2022.0123.

29. Dumitru, C., Kabat, A.M., and Maloy, K.J. (2018). Metabolic Adaptations of CD4(+) T Cells in Inflammatory Disease. Front Immunol 9, 540. 10.3389/fimmu.2018.00540.

30. Luxenburger, H., Thimme, R., and Hofmann, M. (2026). T cell adaptation in chronic infections and tumors. Cell Mol Immunol. 10.1038/s41423-026-01405-y.

31. Nabhan, M., Egan, D., Kreileder, M., Zhernovkov, V., Timosenko, E., Slidel, T., Dovedi, S., Glennon, K., Brennan, D., and Kolch, W. (2023). Deciphering the tumour immune microenvironment cell by cell. Immunooncol Technol 18, 100383. 10.1016/j.iotech.2023.100383.

32. Sanborn, M.A., Wang, X., Gao, S., Dai, Y., and Rehman, J. (2025). Unveiling the cell-type-specific landscape of cellular senescence through single-cell transcriptomics using SenePy. Nat Commun 16, 1884. 10.1038/s41467-025-57047-7.

33. Choo, J., Glisovic, N., and Matic Vignjevic, D. (2022). Gut homeostasis at a glance. J Cell Sci 135. 10.1242/jcs.260248.

34. Ke, G.a.M., Qi and Finley, Thomas and Wang, Taifeng and Chen, Wei and Ma, Weidong and Ye, Qiwei and Liu, Tie-Yan (2017). LightGBM: a highly efficient gradient boosting decision tree. Proceedings of the 31st International Conference on Neural Information Processing Systems. Curran Associates Inc.

35. Wei, Y., Ma, H.K., Wong, M.E., Back, H., Papasavvas, E., Mounzer, K., Aberra, F., Morgenstern, R., Tebas, P., Konnikova, L., et al. (2025). Transcription factor BACH2 shapes tissue-resident memory T cell programs to promote HIV-1 persistence. Immunity 58, 2878–2898 e2811. 10.1016/j.immuni.2025.07.022.

36. Clark, I.C., Mudvari, P., Thaploo, S., Smith, S., Abu-Laban, M., Hamouda, M., Theberge, M., Shah, S., Ko, S.H., Perez, L., et al. (2023). HIV silencing and cell survival signatures in infected T cell reservoirs. Nature 614, 318–325. 10.1038/s41586-022-05556-6.

37. Ventura, J.D., Beloor, J., Allen, E., Zhang, T., Haugh, K.A., Uchil, P.D., Ochsenbauer, C., Kieffer, C., Kumar, P., Hope, T.J., and Mothes, W. (2019). Longitudinal bioluminescent imaging of HIV-1 infection during antiretroviral therapy and treatment interruption in humanized mice. PLoS Pathog 15, e1008161. 10.1371/journal.ppat.1008161.

38. Santangelo, P.J., Rogers, K.A., Zurla, C., Blanchard, E.L., Gumber, S., Strait, K., Connor-Stroud, F., Schuster, D.M., Amancha, P.K., Hong, J.J., et al. (2015). Whole-body immunoPET reveals active SIV dynamics in viremic and antiretroviral therapy-treated macaques. Nat Methods 12, 427–432. 10.1038/nmeth.3320.

39. Samer, S., Thomas, Y., Arainga, M., Carter, C., Shirreff, L.M., Arif, M.S., Avita, J.M., Frank, I., McRaven, M.D., Thuruthiyil, C.T., et al. (2023). Blockade of TGF-beta signaling reactivates HIV-1/SIV reservoirs and immune responses in vivo. JCI Insight 8. 10.1172/jci.insight.176882.

40. Taylor, R.A., Xiao, S., Carias, A.M., McRaven, M.D., Thakkar, D.N., Arainga, M., Allen, E.J., Rogers, K.A., Kumarapperuma, S.C., Gong, S., et al. (2021). PET/CT targeted tissue sampling reveals virus specific dIgA can alter the distribution and localization of HIV after rectal exposure. PLoS Pathog 17, e1009632. 10.1371/journal.ppat.1009632.

41. Sanders-Beer, B.E., Eschricht, M., Seifried, J., Hirsch, V.M., Allan, J.S., and Norley, S. (2012). Characterization of a monoclonal anti-capsid antibody that cross-reacts with three major primate lentivirus lineages. Virology 422, 402–412. 10.1016/j.virol.2011.11.003.

42. Hao, Y., Stuart, T., Kowalski, M.H., Choudhary, S., Hoffman, P., Hartman, A., Srivastava, A., Molla, G., Madad, S., Fernandez-Granda, C., and Satija, R. (2024). Dictionary learning for integrative, multimodal and scalable single-cell analysis. Nat Biotechnol 42, 293–304. 10.1038/s41587-023-01767-y.

43. Korsunsky, I., Millard, N., Fan, J., Slowikowski, K., Zhang, F., Wei, K., Baglaenko, Y., Brenner, M., Loh, P.R., and Raychaudhuri, S. (2019). Fast, sensitive and accurate integration of single-cell data with Harmony. Nat Methods 16, 1289–1296. 10.1038/s41592-019-0619-0.

44. Finak, G., McDavid, A., Yajima, M., Deng, J., Gersuk, V., Shalek, A.K., Slichter, C.K., Miller, H.W., McElrath, M.J., Prlic, M., et al. (2015). MAST: a flexible statistical framework for assessing transcriptional changes and characterizing heterogeneity in single-cell RNA sequencing data. Genome Biol 16, 278. 10.1186/s13059-015-0844-5.

45. Kueckelhaus, J., Frerich, S., Kada-Benotmane, J., Koupourtidou, C., Ninkovic, J., Dichgans, M., Beck, J., Schnell, O., and Heiland, D.H. (2024). Inferring histology-associated gene expression gradients in spatial transcriptomic studies. Nat Commun 15, 7280. 10.1038/s41467-024-50904-x.

46. Subramanian, A., Tamayo, P., Mootha, V.K., Mukherjee, S., Ebert, B.L., Gillette, M.A., Paulovich, A., Pomeroy, S.L., Golub, T.R., Lander, E.S., and Mesirov, J.P. (2005). Gene set enrichment analysis: a knowledge-based approach for interpreting genome-wide expression profiles. Proc Natl Acad Sci U S A 102, 15545–15550. 10.1073/pnas.0506580102.

47. Holland, P.B.-i.-M.a.J.V.S.a.J.B.a.C.G.a.D.D.a.S.M.-D.a.P.T.a.A.D.a.C.H. (2022). decoupleR: Ensemble of computational methods to infer biological activities from omics data.. Bioinformatics Advances. 10.1093/bioadv/vbac016.

48. Lopez, R., Li, B., Keren-Shaul, H., Boyeau, P., Kedmi, M., Pilzer, D., Jelinski, A., Yofe, I., David, E., Wagner, A., et al. (2022). DestVI identifies continuums of cell types in spatial transcriptomics data. Nat Biotechnol 40, 1360–1369. 10.1038/s41587-022-01272-8.

49. Han, L., Wei, X., Liu, C., Volpe, G., Zhuang, Z., Zou, X., Wang, Z., Pan, T., Yuan, Y., Zhang, X., et al. (2022). Cell transcriptomic atlas of the non-human primate Macaca fascicularis. Nature 604, 723–731. 10.1038/s41586-022-04587-3.

50. Elmentaite, R., Kumasaka, N., Roberts, K., Fleming, A., Dann, E., King, H.W., Kleshchevnikov, V., Dabrowska, M., Pritchard, S., Bolt, L., et al. (2021). Cells of the human intestinal tract mapped across space and time. Nature 597, 250–255. 10.1038/s41586-021-03852-1.

51. Wolf, F.A., Angerer, P., and Theis, F.J. (2018). SCANPY: large-scale single-cell gene expression data analysis. Genome Biol 19, 15. 10.1186/s13059-017-1382-0.

52. James, K.R., Gomes, T., Elmentaite, R., Kumar, N., Gulliver, E.L., King, H.W., Stares, M.D., Bareham, B.R., Ferdinand, J.R., Petrova, V.N., et al. (2020). Distinct microbial and immune niches of the human colon. Nat Immunol 21, 343–353. 10.1038/s41590-020-0602-z.

53. Pham, D., Tan, X., Balderson, B., Xu, J., Grice, L.F., Yoon, S., Willis, E.F., Tran, M., Lam, P.Y., Raghubar, A., et al. (2023). Robust mapping of spatiotemporal trajectories and cell-cell interactions in healthy and diseased tissues. Nat Commun 14, 7739. 10.1038/s41467-023-43120-6.

54. Bischl, M.L.a.M.B.a.J.R.a.P.S.a.F.P.a.S.C.a.Q.A.a.G.C.a.L.K.a.B. (2019). mlr3: A modern object-oriented machine learning framework in R. Journal of Open Source Software. 10.21105/joss.01903.

