## Supplementary figures and images for "A Tissue Virus Microenvironment with Activated Stress Responses Underlies Durable SIV Persistence"

### Supplementary Figure 1

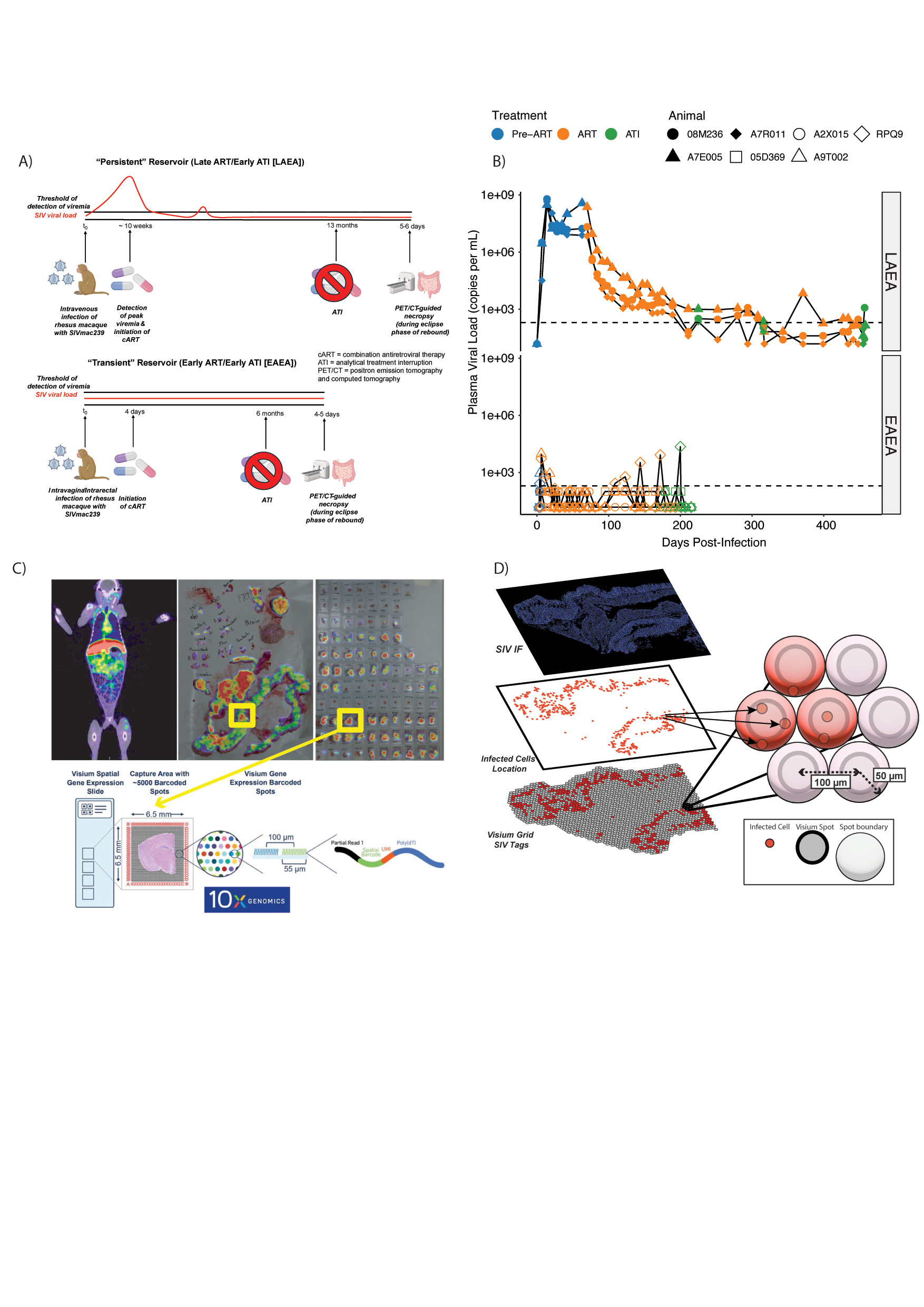

### Supplementary Figure 2

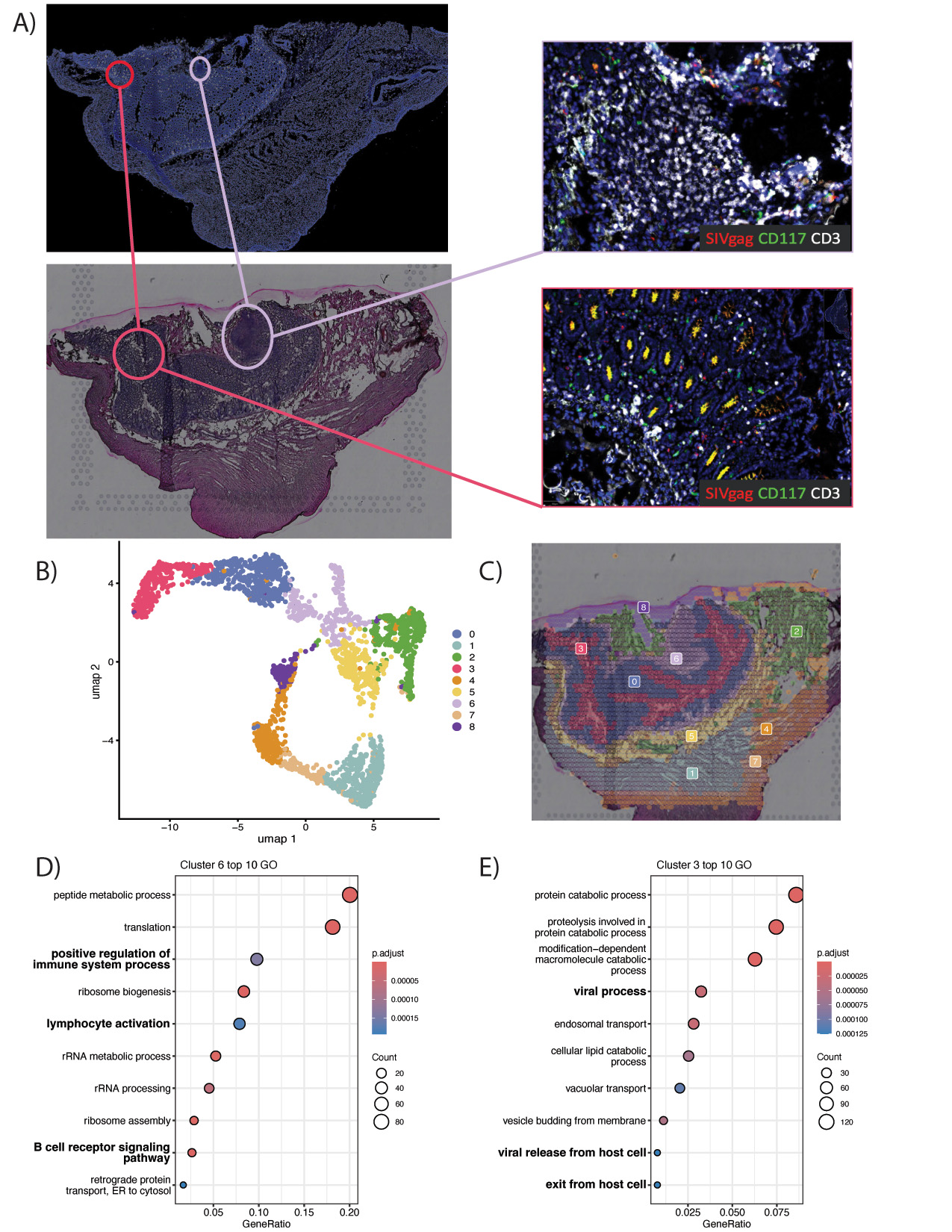

### Supplementary Figure 3

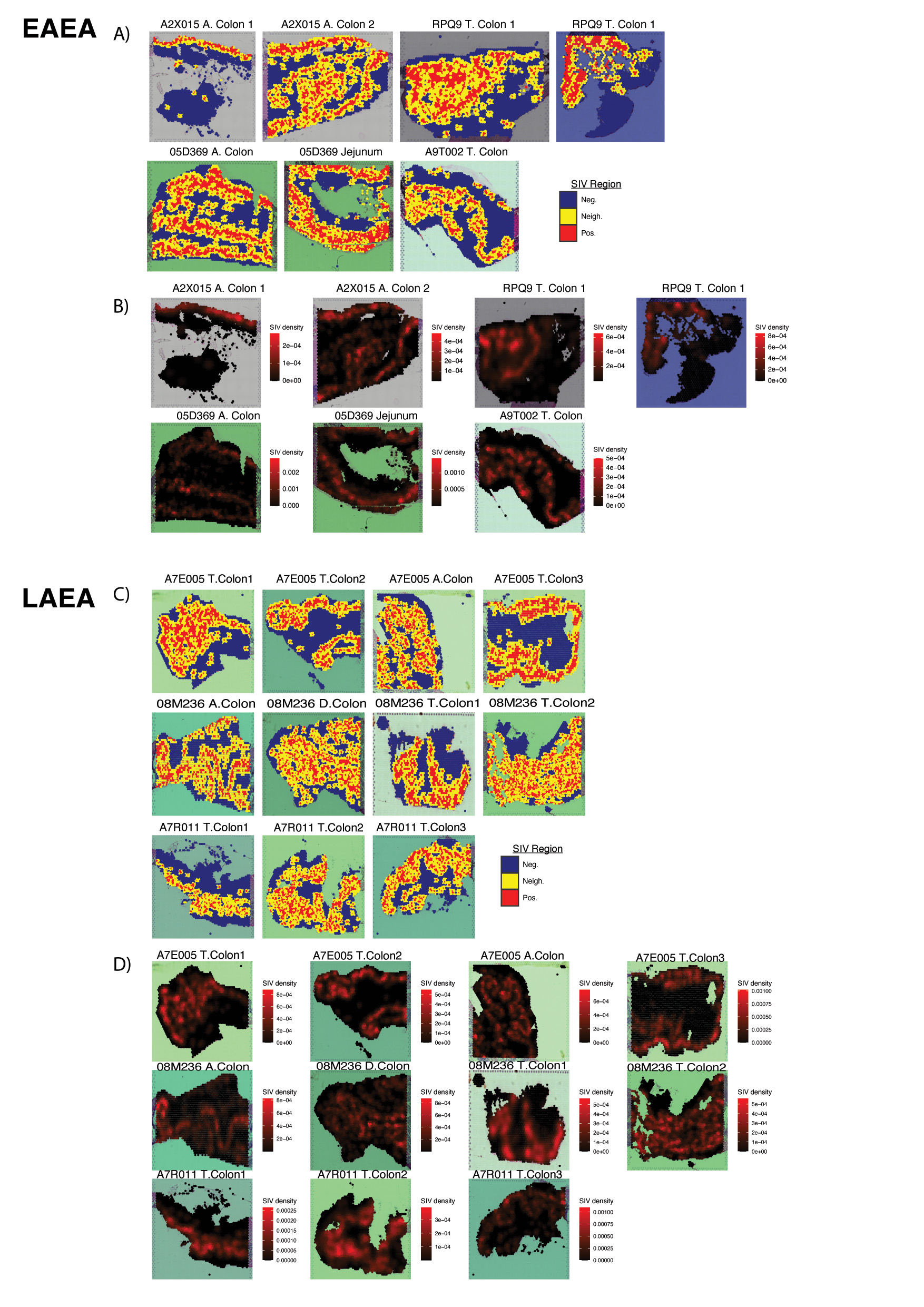

### Supplementary Figure 4

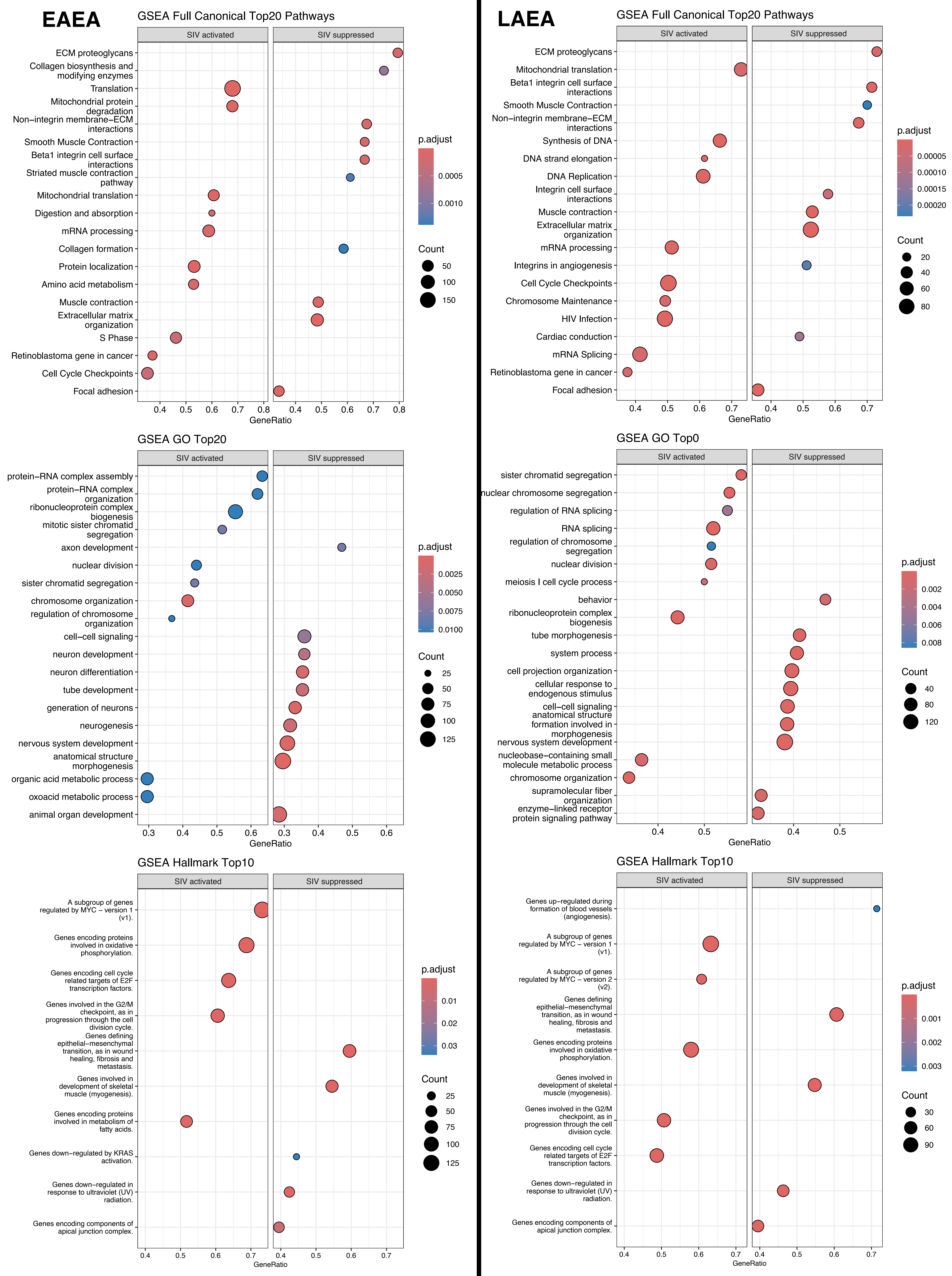

### Supplementary Figure 5

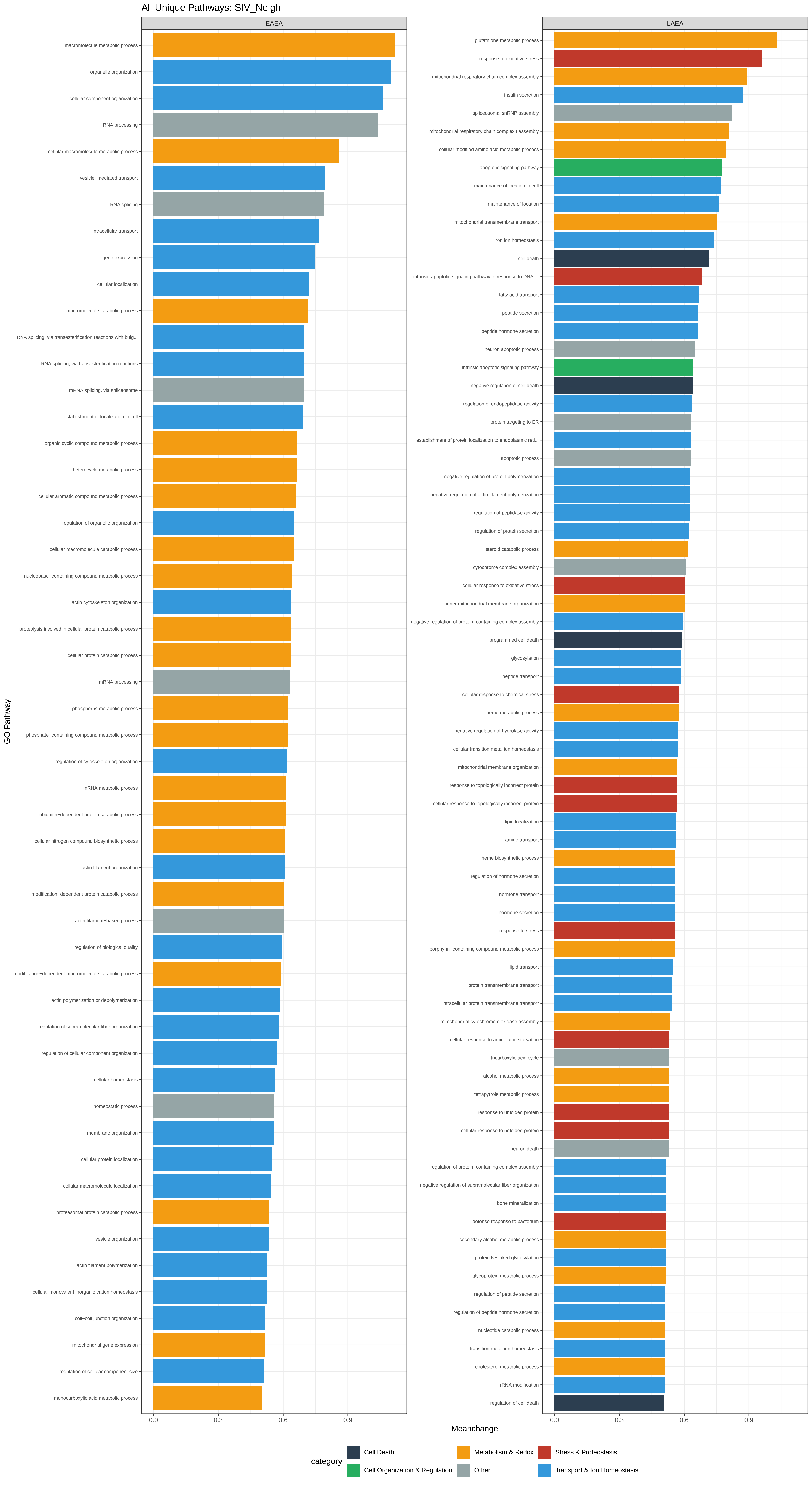

### Supplementary Figure 6

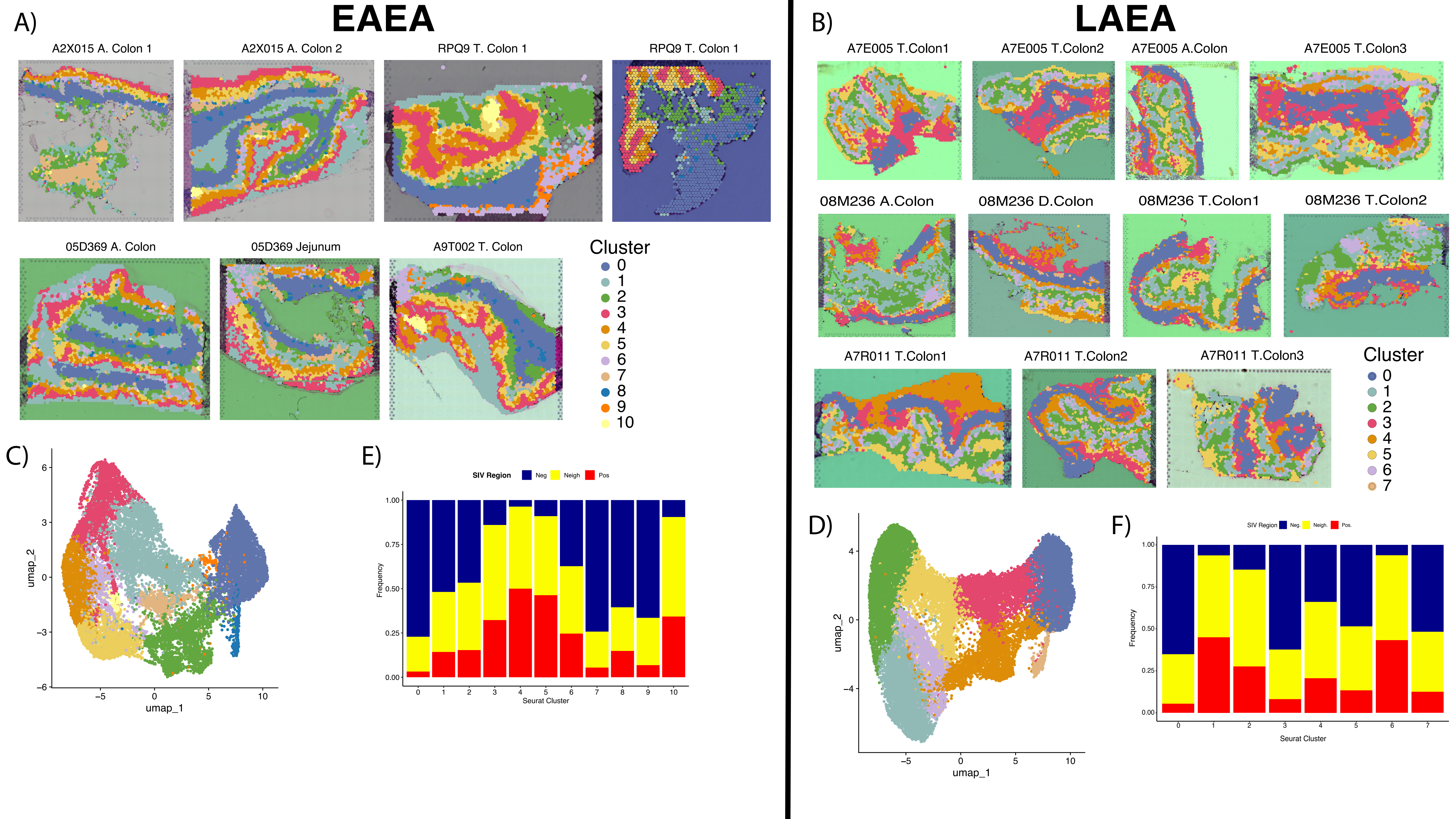

### Supplementary Figure 7

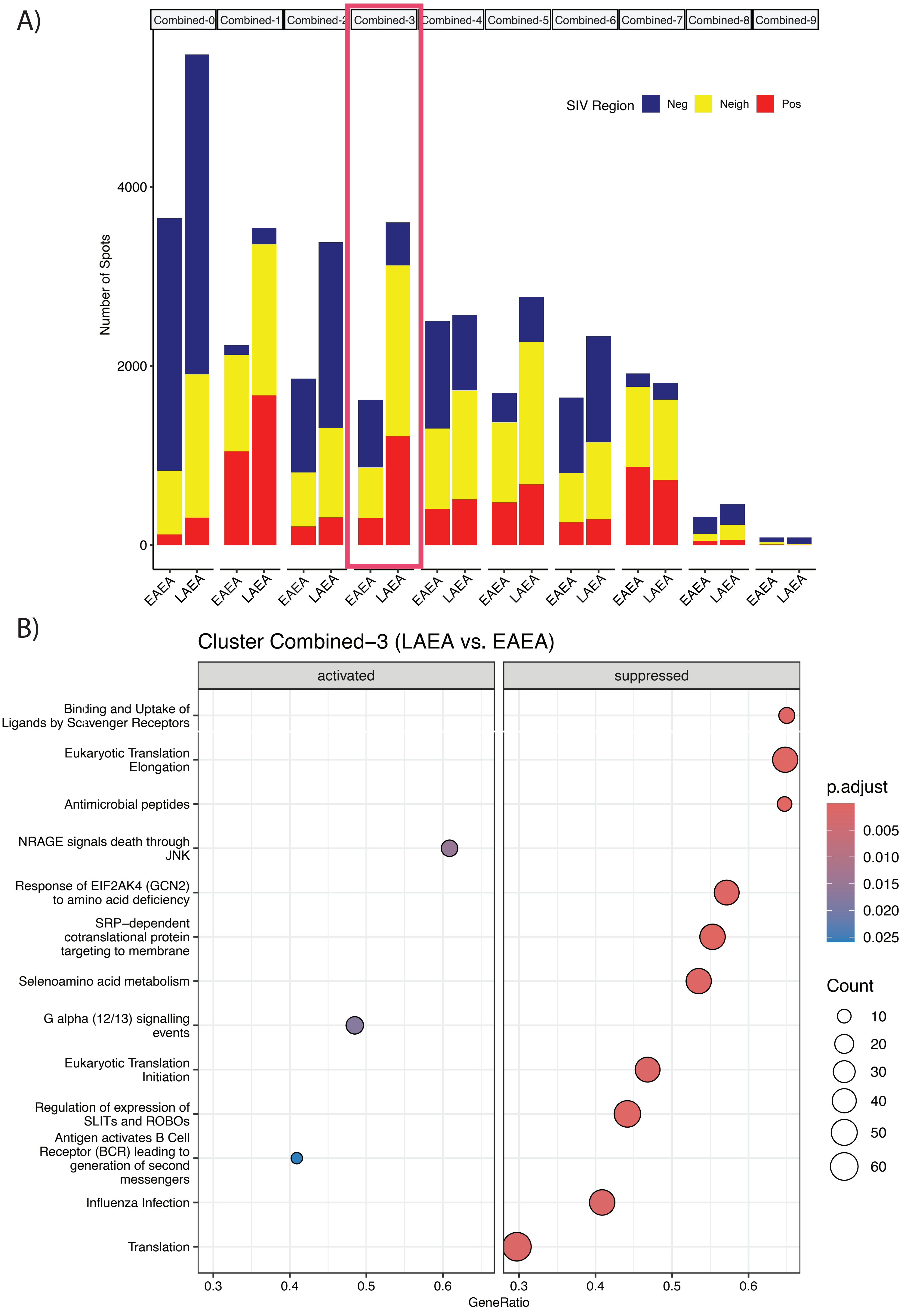

### Supplementary Figure 8

#### EAEA slide
(Broad gut cells)

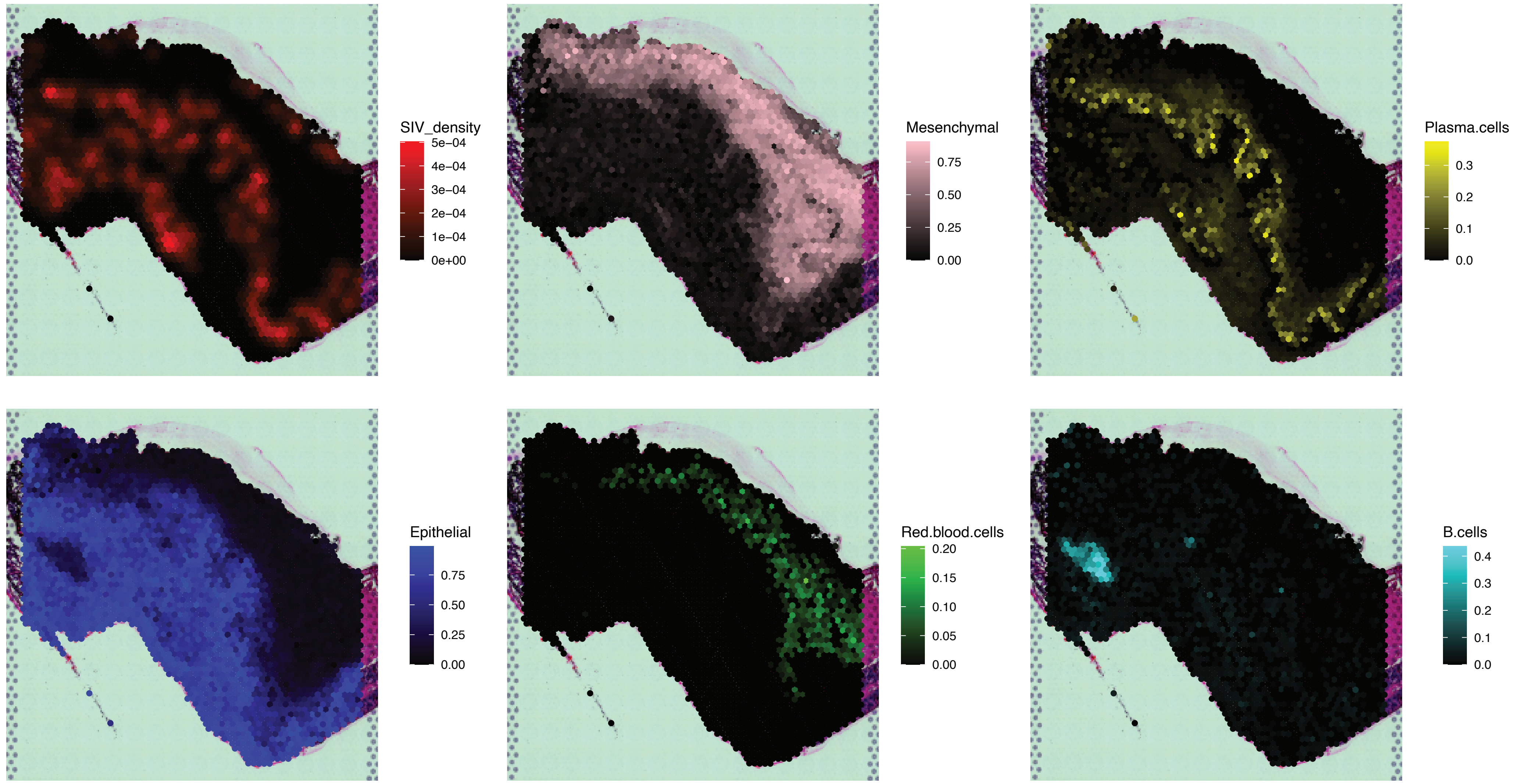


#### LAEA slide
(Broad gut cells)

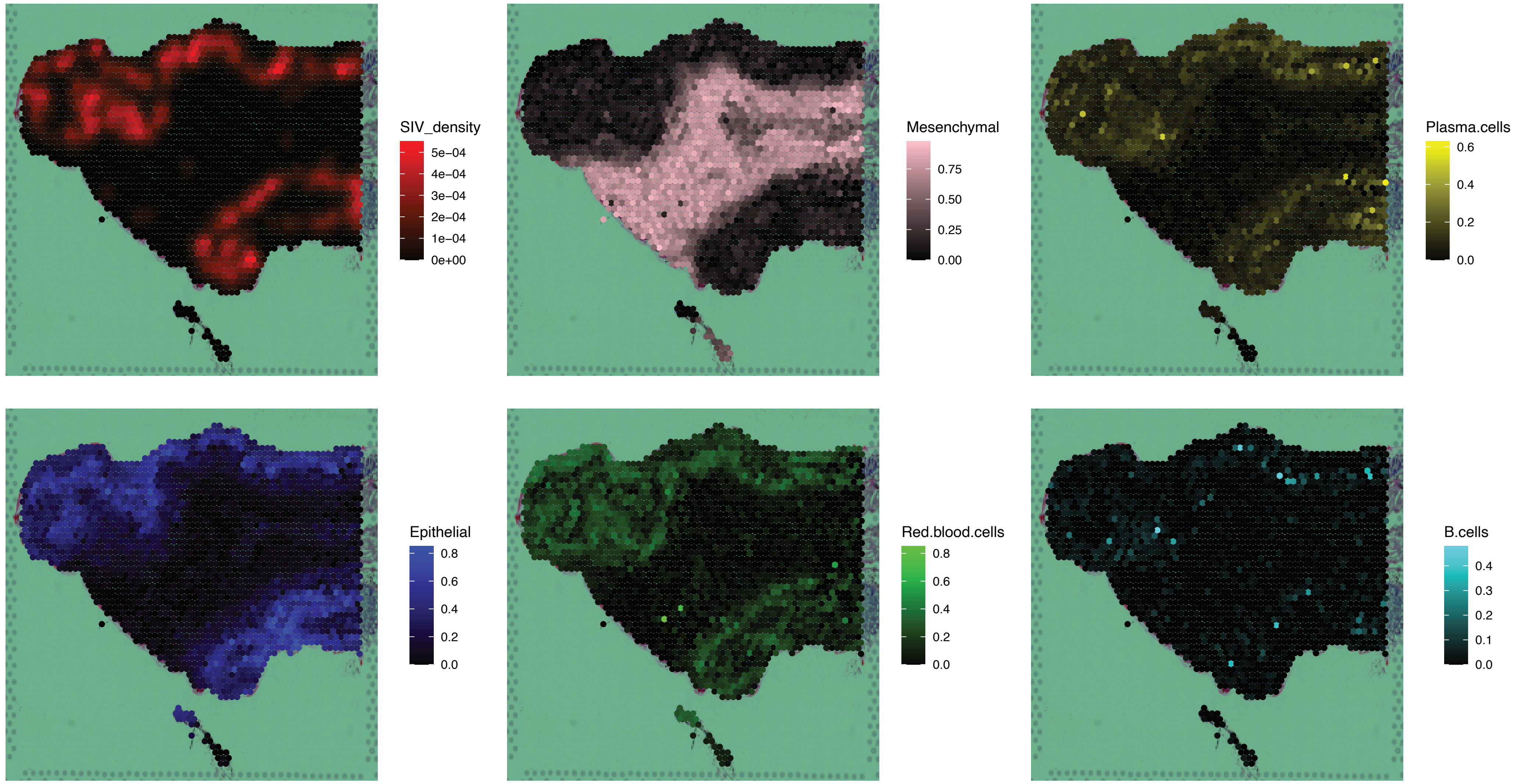

### Supplementary Figure 9

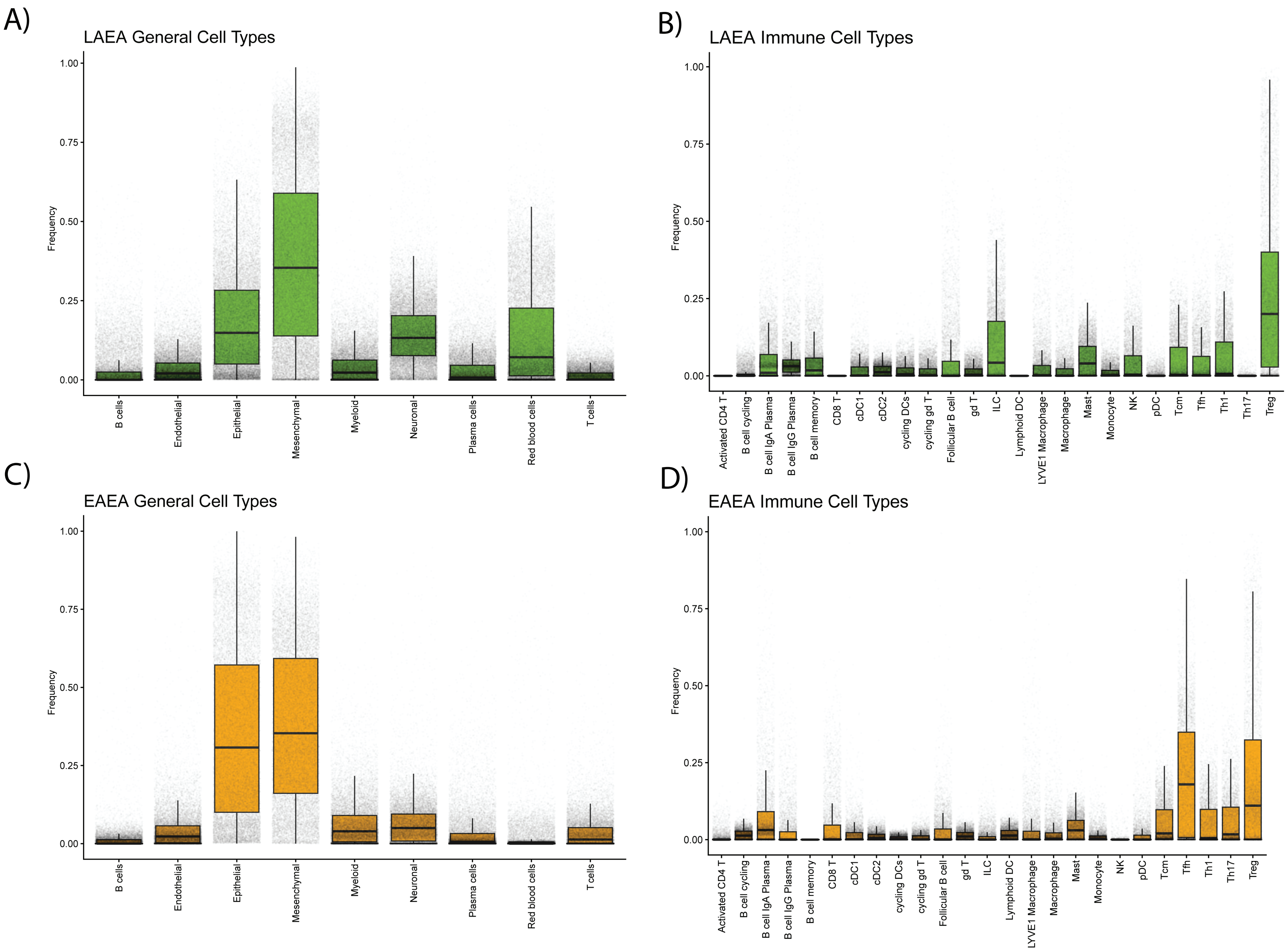

### Supplementary Figure 10

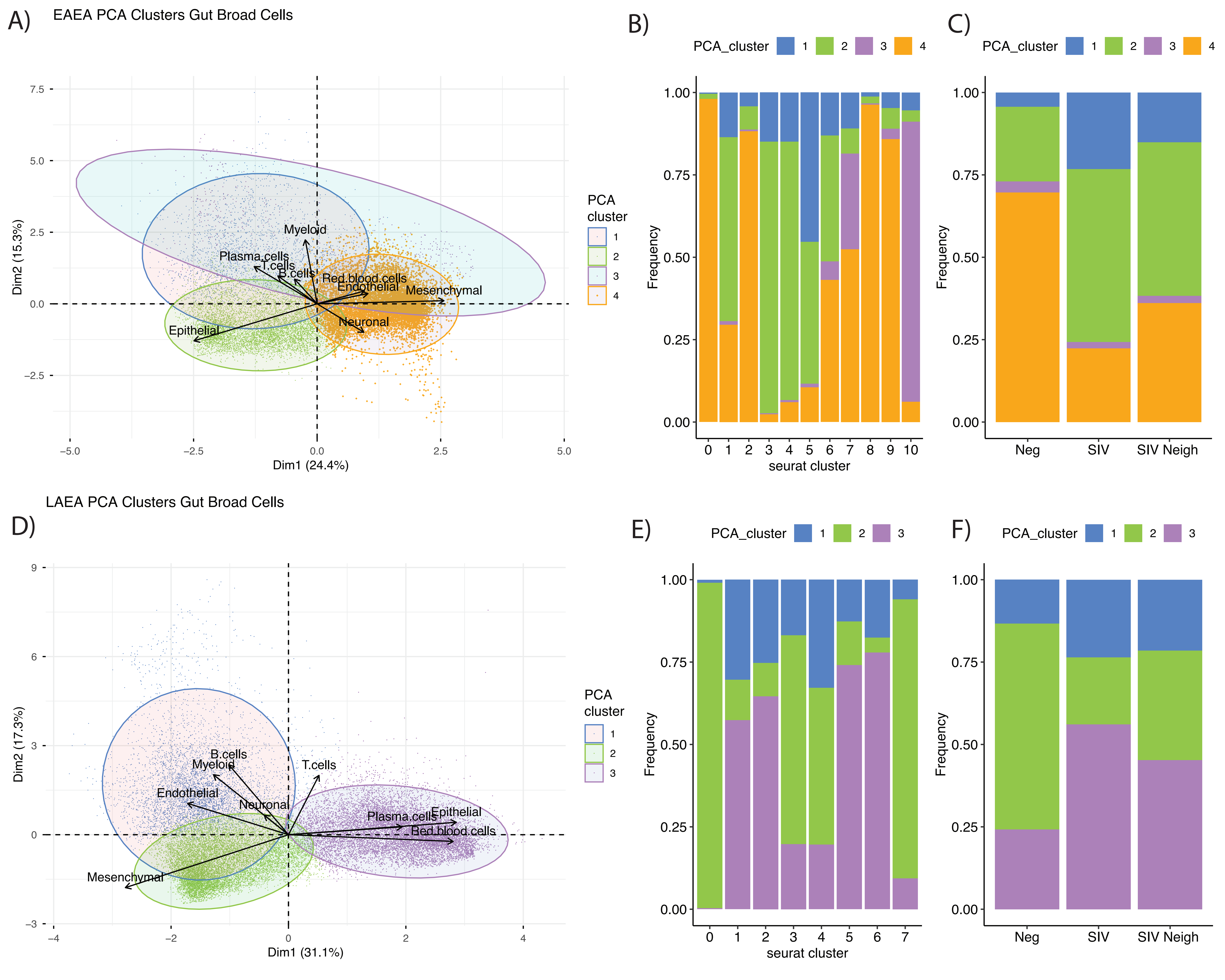

### Supplementary Figure 11

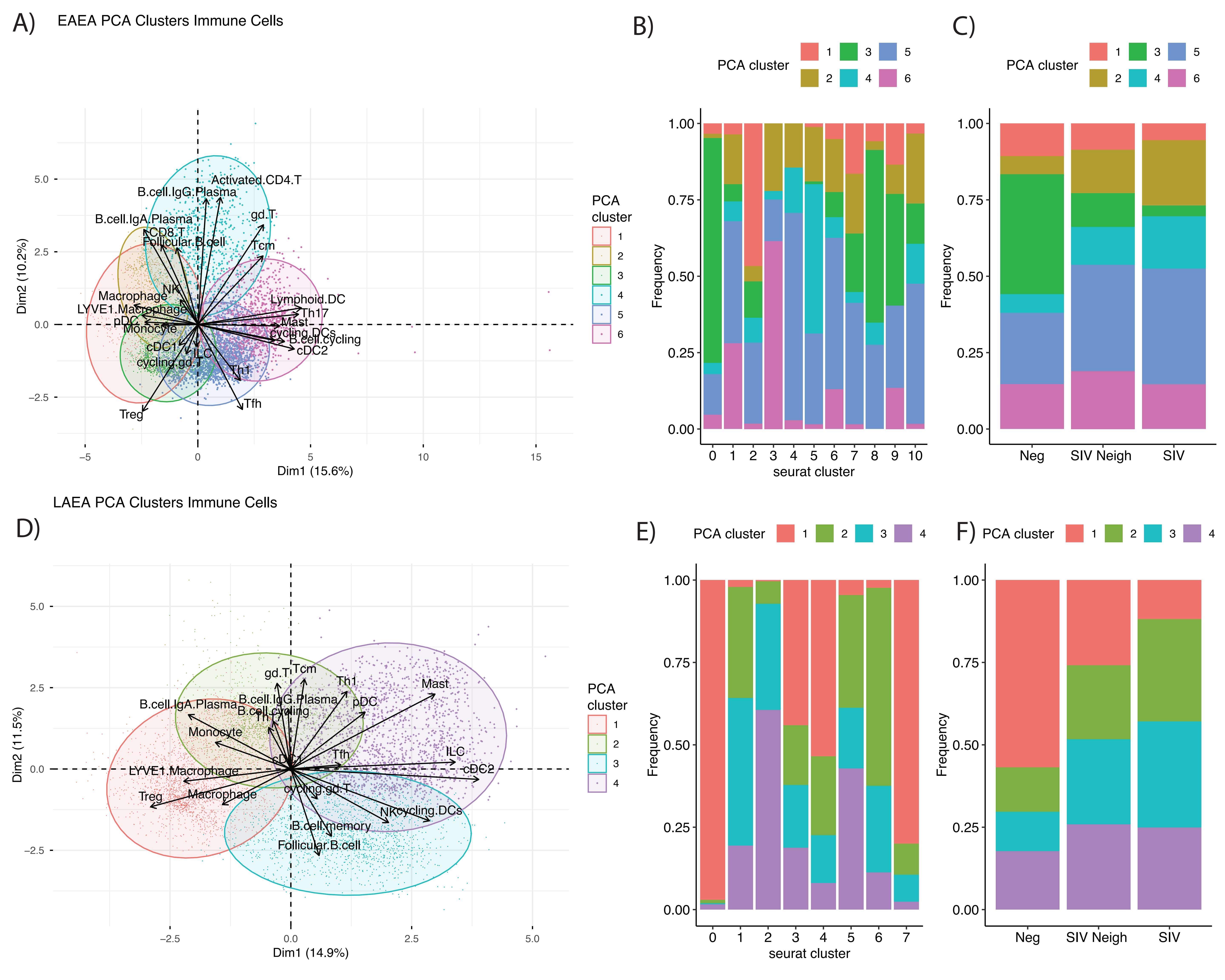

### Supplementary Figure 12

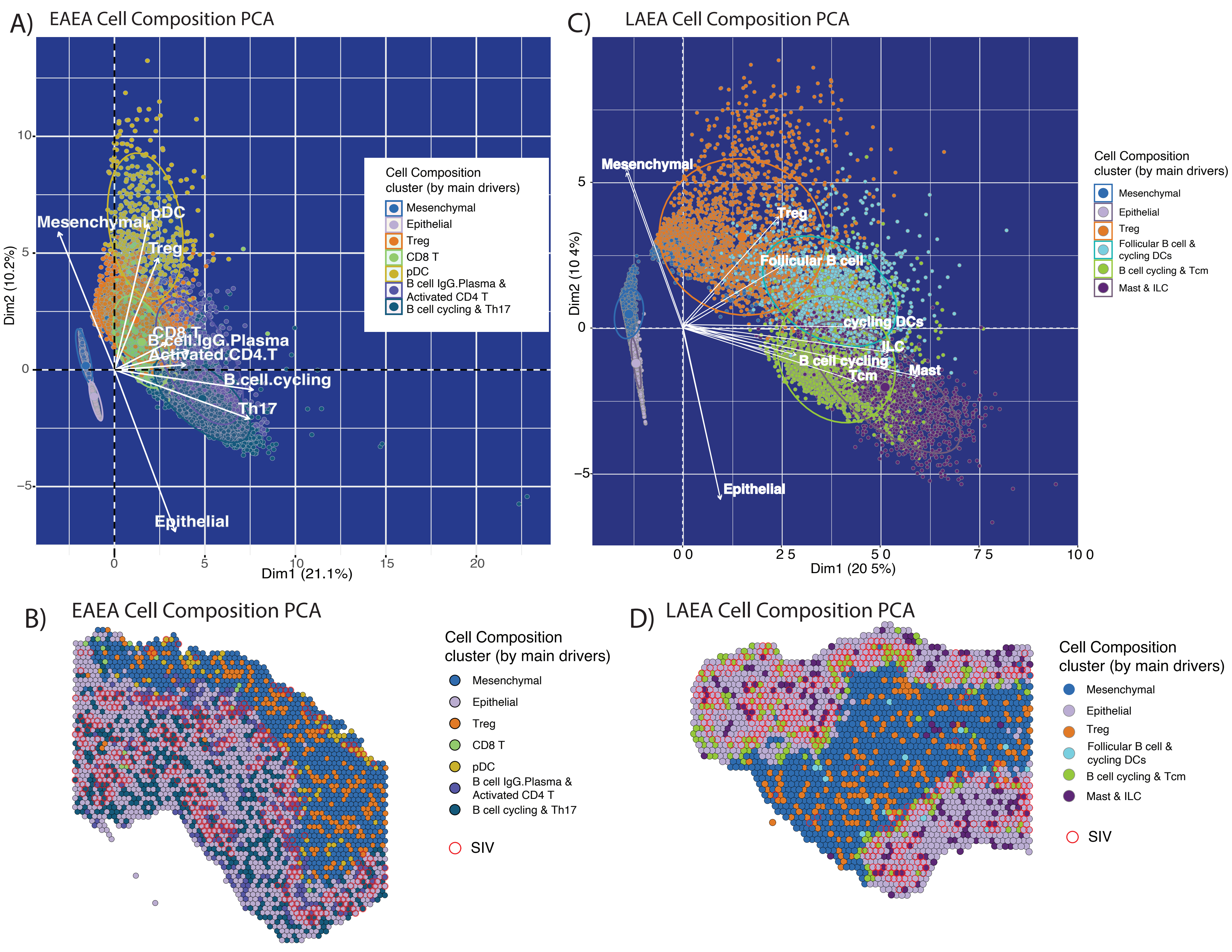

### Supplementary Figure 13

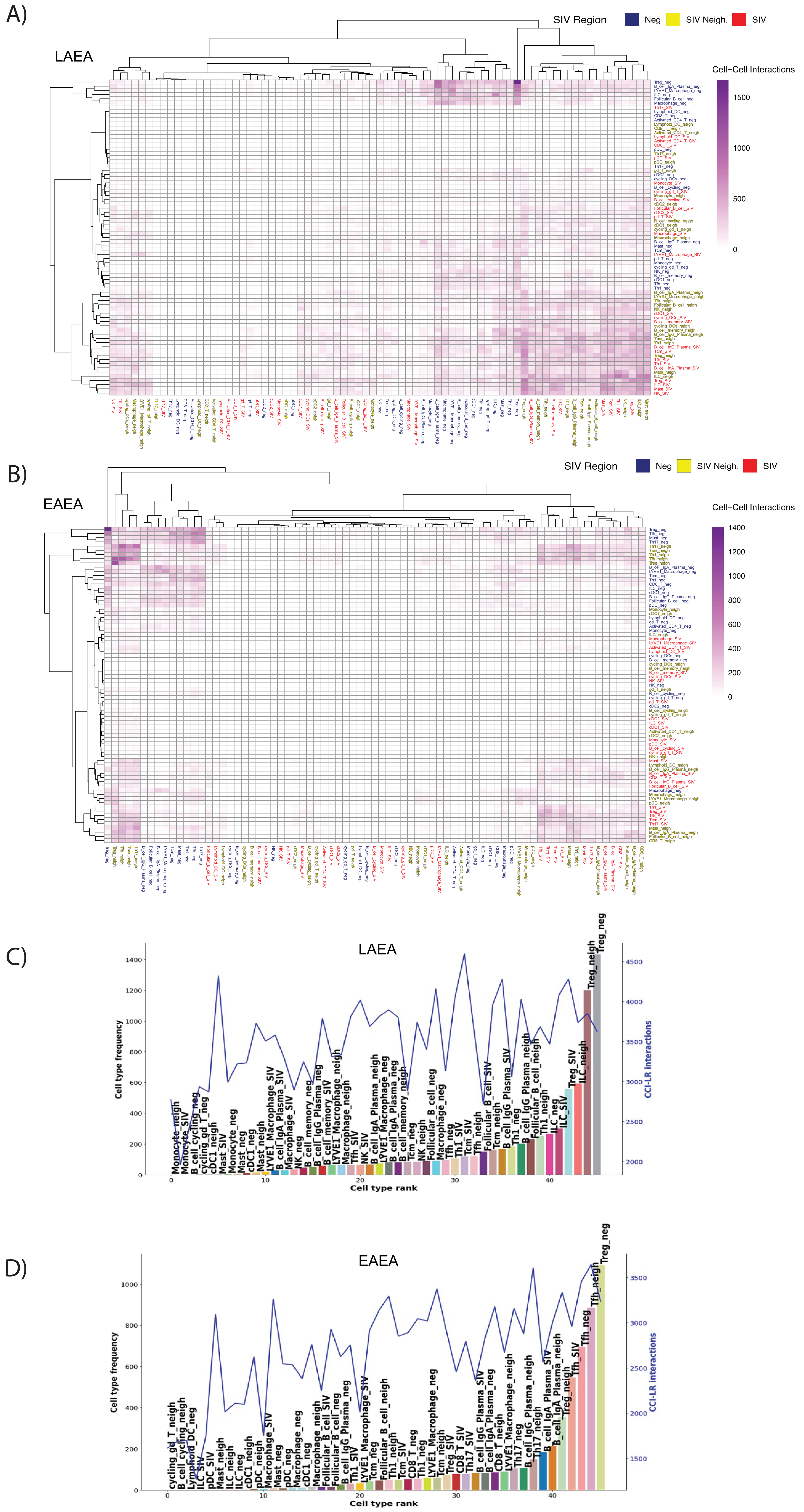

### Supplementary Figure 14

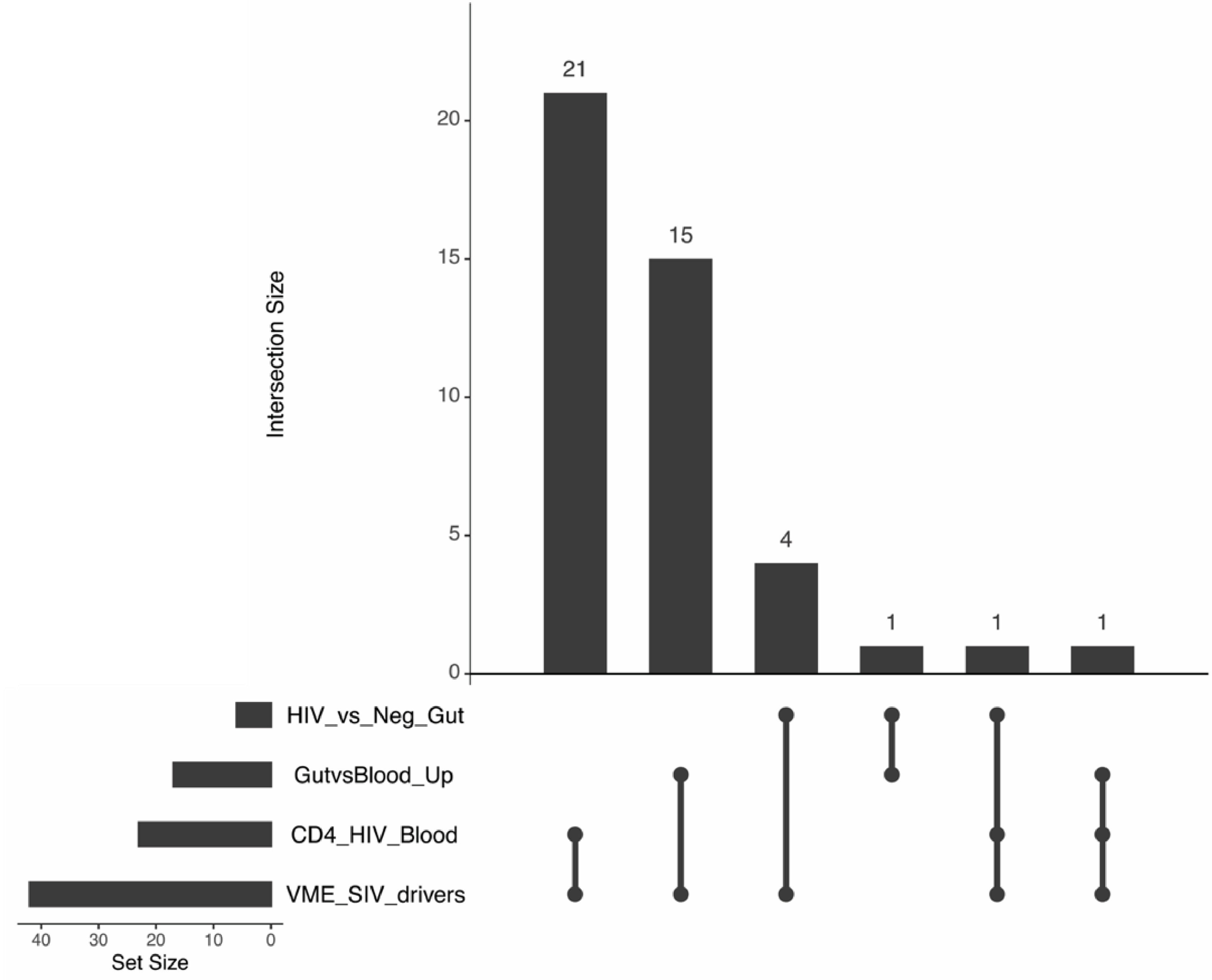

### Supplementary Figure 15

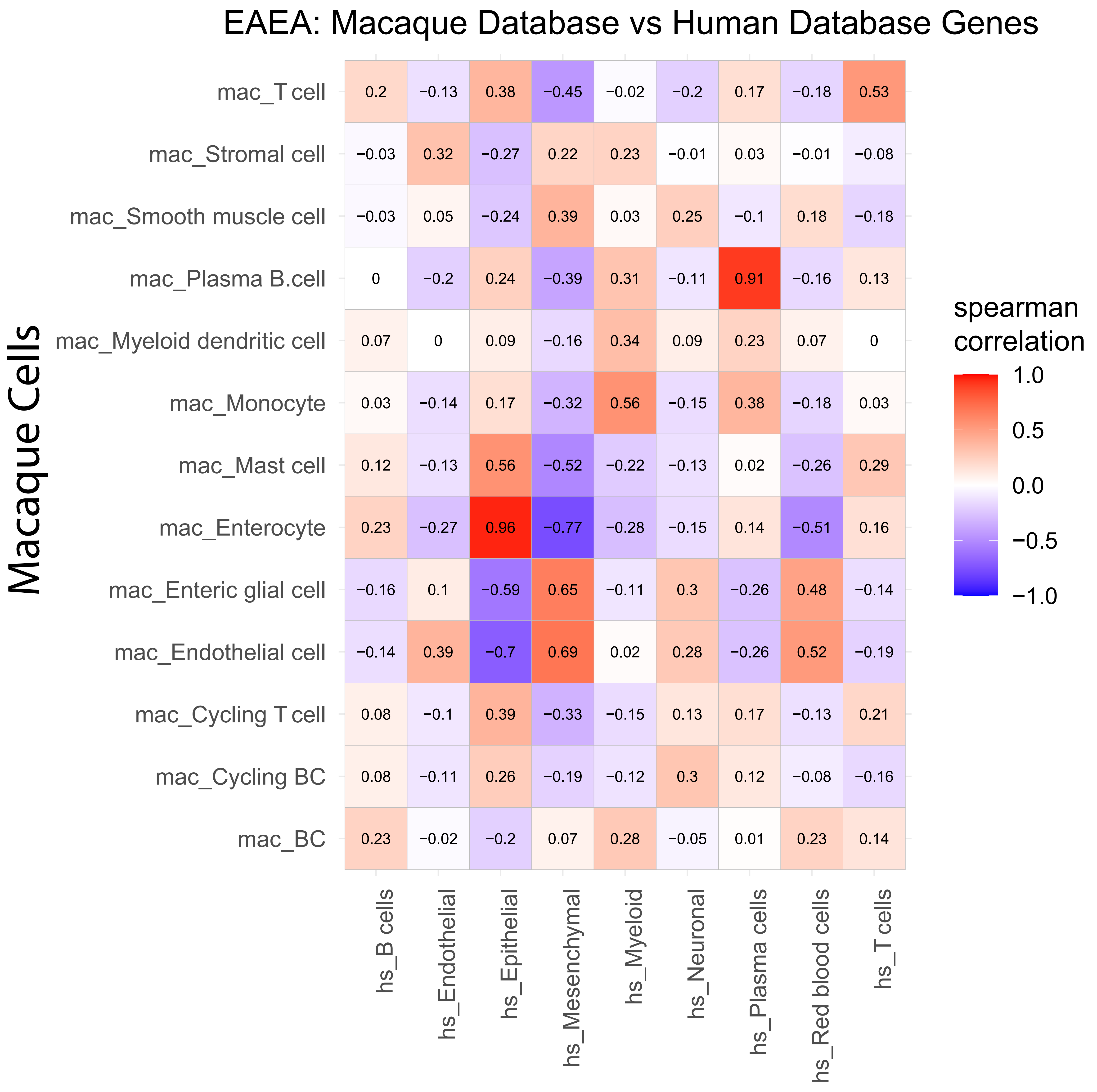

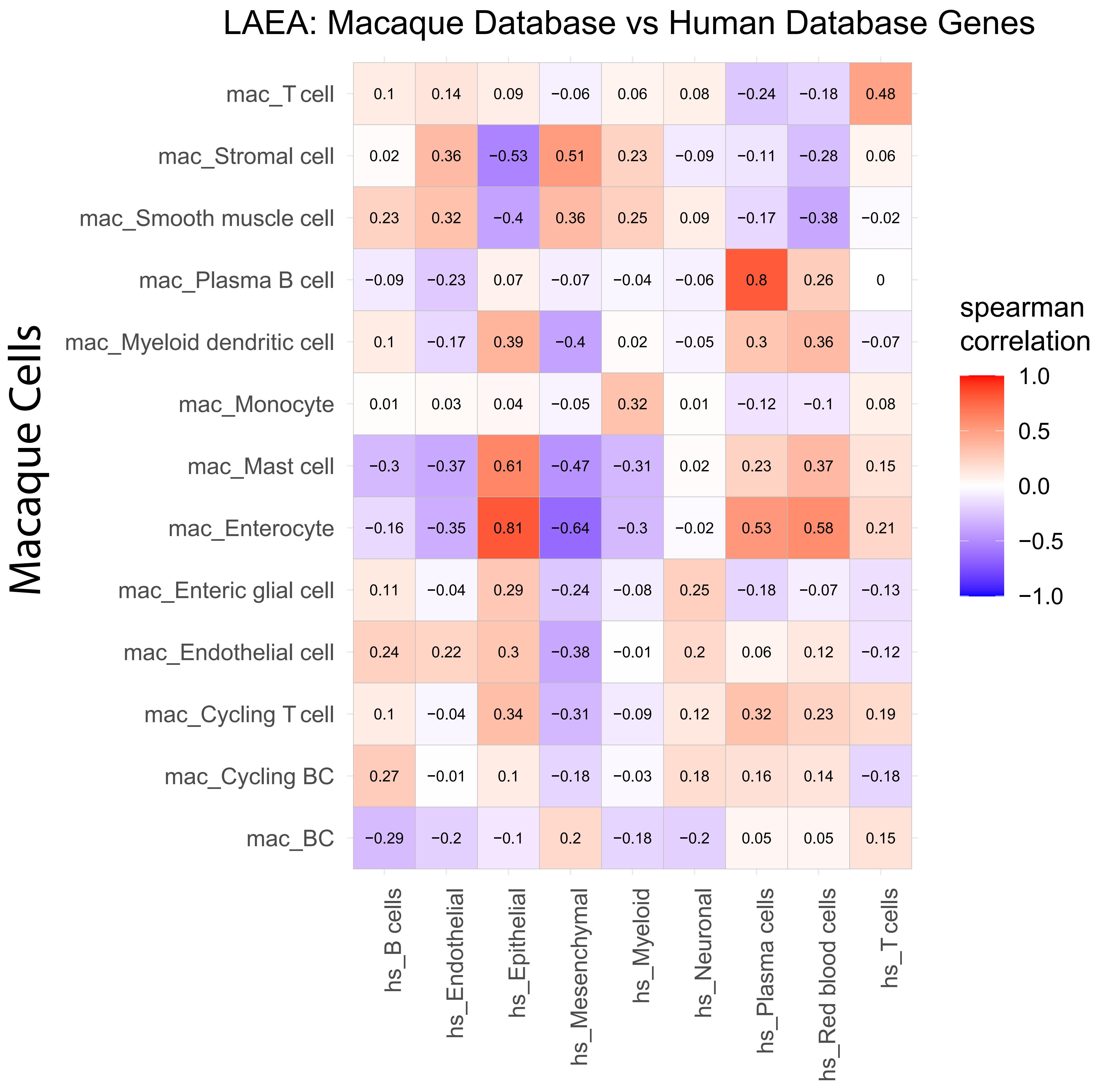
