## Supplementary Table 3 for "A Tissue Virus Microenvironment with Activated Stress Responses Underlies Durable SIV Persistence"

| **Animal** | **Sex** | **Infection** | **Viral challenge** | **ART START** | **ART END** | **Necropsy** | **Treatment (days)** | **Treatment (months)** | **Days post ATI** |
| --- | --- | --- | --- | --- | --- | --- | --- | --- | --- |
| A9T002 | F | SIVmac239 | 17-Aug-19 | 21-Aug-19 | 9-Feb-20 | 13-Feb-20 | 172 | 5 | 4 |
| RPQ9 | F | SIVmac239 | 17-Aug-19 | 21-Aug-19 | 1-Mar-20 | 5-Mar-20 | 193 | 6 | 4 |
| A2X015 | F | SIVmac239 | 18-Nov-18 | 21-Nov-18 | 14-Apr-19 | 19-Apr-19 | 144 | 4 | 5 |
| 05D369 | F | SIV-ffLuc | 19-Jan-19 | 23-Jan-19 | 14-Jul-19 | 8-Aug-09 | 172 | 5 | 4 |
| A7E005 | M | SIVmac239M2 | 20-Jul-21 | 28-Sep-21 | 15-Oct-22 | 21-Oct-22 | 382 | 12 | 6 |
| A7R011 | F | SIVmac239M2 | 20-Jul-21 | 28-Sep-21 | 15-Oct-22 | 20-Oct-22 | 382 | 12 | 5 |
| 08M236 | F | SIVmac239M2 | 20-Jul-21 | 28-Sep-21 | 15-Oct-22 | 20-Oct-22 | 382 | 12 | 5 |

**Supplementary Table 3**
