## Supplementary Table 4 for "A Tissue Virus Microenvironment with Activated Stress Responses Underlies Durable SIV Persistence"

| **Atlas** | **Cell_ID** | **Full_Name** |
| --- | --- | --- |
| Pan-GI Gut Cell Atlas | B.cells | B Cells |
| Pan-GI Gut Cell Atlas | Endothelial | Endothelial Cells |
| Pan-GI Gut Cell Atlas | Epithelial | Epithelial Cells |
| Pan-GI Gut Cell Atlas | Mesenchymal | Mesenchymal Cells |
| Pan-GI Gut Cell Atlas | Myeloid | Myeloid Cells |
| Pan-GI Gut Cell Atlas | Neuronal | Neuronal Cells |
| Pan-GI Gut Cell Atlas | Plasma.cells | Plasma Cells |
| Pan-GI Gut Cell Atlas | Red.blood.cells | Red Blood Cells |
| Pan-GI Gut Cell Atlas | T.cells | T Cells |
| Colon Immune Cell Atlas | Activated.CD4.T | Activated CD4+ T Cells |
| Colon Immune Cell Atlas | B.cell.IgA.Plasma | IgA Plasma B Cells |
| Colon Immune Cell Atlas | B.cell.IgG.Plasma | IgG Plasma B Cells |
| Colon Immune Cell Atlas | B.cell.cycling | Cycling B Cells |
| Colon Immune Cell Atlas | Follicular.B.cell | Follicular B Cells |
| Colon Immune Cell Atlas | B.cell.memory | Memory B Cells |
| Colon Immune Cell Atlas | CD8.T | CD8+ T Cells |
| Colon Immune Cell Atlas | ILC | Innate Lymphoid Cells |
| Colon Immune Cell Atlas | Lymphoid.DC | Lymphoid Dendritic Cells |
| Colon Immune Cell Atlas | Monocyte | Monocytes |
| Colon Immune Cell Atlas | Mast | Mast Cells |
| Colon Immune Cell Atlas | Macrophage | Macrophages |
| Colon Immune Cell Atlas | LYVE1.Macrophage | LYVE1+ Macrophages |
| Colon Immune Cell Atlas | NK | Natural Killer Cells |
| Colon Immune Cell Atlas | Tcm | Central Memory T Cells |
| Colon Immune Cell Atlas | Tfh | T Follicular Helper Cells |
| Colon Immune Cell Atlas | Th1 | T Helper 1 Cells |
| Colon Immune Cell Atlas | Th17 | T Helper 17 Cells |
| Colon Immune Cell Atlas | Treg | Regulatory T Cells |
| Colon Immune Cell Atlas | cDC1 | Conventional Dendritic Cells Type 1 |
| Colon Immune Cell Atlas | cDC2 | Conventional Dendritic Cells Type 2 |
| Colon Immune Cell Atlas | cycling.DCs | Cycling Dendritic Cells |
| Colon Immune Cell Atlas | pDC | Plasmacytoid Dendritic Cells |
| Colon Immune Cell Atlas | gd.T | Gamma-Delta T Cells |
| Colon Immune Cell Atlas | cycling.gd.T | Cycling Gamma-Delta T Cells |
